# Single-Cell Profiling Reveals Developmental Trajectories and identifies SYK and TIM3 as Targets in some T Cell Lymphomas

**DOI:** 10.64898/2026.03.27.714741

**Authors:** Ruoyan Li, Jamie D. Matthews, Emily R. James, Carla Vazquez-Amos, Olli Dufva, Shuang Li, Christopher J. Steel, Lorenz Kretschmer, Chenqu Suo, Philip Turton, Ruth F. Jarrett, Ekaterina Shelomentseva, Egor Volchkov, Dmitry Abramov, Maria Myrsini Tzioni, Ming-Qing Du, Olaf Merkel, Michaela Schlederer, Lukas Kenner, Sarah A. Teichmann, Suzanne D. Turner

## Abstract

T cell lymphomas (TCL) are a heterogeneous collection of malignancies whose origins and pathogenesis are poorly understood and for which few efficacious therapeutic options exist. Here, we conduct single-cell transcriptomic profiling spanning eight TCL entities and describe entity-associated programmes. We predict the cell of origin for these tumours through an integrative analysis of transcriptome and T cell receptor (TCR) maturation states. By identifying tumours with TCR states ranging from the pre-TCR through non-productive and productive TCR alpha and beta chain rearrangements we shed new light on their developmental origins. Furthermore, we apply our drug2cell computational drug target predictions with drug screens using patient-derived cell models, systematically benchmarking the performance of drug2cell and validating compounds and targets. This process identifies SYK inhibitors as a therapeutic opportunity and prioritises TIM3 for immunotherapy based on combined spatial transcriptomics analysis. Overall, our data provide a resource for diagnostics and therapies for tumours of critical unmet need.

## Introduction

T cell lymphomas (TCLs) are a heterogenous class of malignancies originating in the immune system.^1^ Largely, they are defined as deriving from either immature or mature T cells with involvement of nodal and/or extranodal sites. Sub-classification can be difficult; careful histopathology, combined with genomics/genetics and clinical information on disease presentation are required for diagnosis.^1–3^ However, disease heterogeneity and ongoing studies ensure an evolving landscape, most recently exemplified by the reclassification of nodal T Follicular Helper lymphomas (nTFHL) and continued relegation of some cases by diagnostic exclusion to the ‘wastebasket’ category of peripheral T cell lymphoma, not otherwise specified (PTCL-NOS).^1^

The molecular genetics underlying these malignancies are equally heterogeneous, although some oncogenic drivers are known, notably Nucleophosmin 1-Anaplastic Lymphoma Kinase (NPM1-ALK), aberrantly expressed as the result of a t(2;5)(p23;q35) in ALK+ anaplastic large cell lymphoma (ALCL) which can also be defined by other ALK fusion partners.^4,5^ Investigations into ALK-negative ALCL have identified single nucleotide variants and chromosomal rearrangements activating the JAK/STAT pathway, which has also been implicated in the pathogenesis of breast implant-associated ALCL (BIA-ALCL)^6–10^, now a distinct clinical entity in the recent WHO classification of haematolymphoid tumours.^1^ Our knowledge of T lymphoblastic lymphoma (TLBL) has also improved through genomic characterisation revealing prognostic associations.^11^ In comparison, PTCL-NOS has been slow to follow, although some defining genomic features have led to sub-classification into PTCL-GATA3 or PTCL-TBX21 (with or without DNMT3A mutations).^9,12,13^

Whilst genomic discoveries have led to improved therapeutic approaches for some TCLs, for others, little has changed over the years. For example, outcomes for patients diagnosed with ALK^+^ ALCL treated according to the ‘ALCL99’ multi-agent chemotherapy protocol have a 10-year overall survival of 90%^14^ which has improved with the introduction of ALK inhibitors and brentuximab.^15,16^ In contrast, treatment regimens and outcomes have changed little for PTCL-NOS, with CHOP (cyclophosphamide, doxorubicin, vincristine, and prednisone) being a mainstay with complete response rates of just 30%.^17^ In order to investigate these issues further, we conducted single-cell RNA sequencing (scRNA-seq) and *T cell receptor* (*TR*) sequencing to create a broad TCL dataset, revealing new insights into cell of origin classifications and potential mechanisms of pathogenesis. Furthermore, we predict novel oncogenic mutations and drug targets, and validate them with systematic *in vitro* screens of PDX tissue, showing consistency between *in silico* and *in vitro* approaches.

## Methods

### Patient samples

Aliquots of fresh-frozen disaggregated TLBL and PTCL were obtained from the Glasgow viable lymphoma cell bank under research ethics committee reference number 07/Q0104/16 approved by the Huntingdon research ethics committee, Cambridge, UK. ALCL patient derived xenografts (PDX) were developed from fresh disaggregated primary patient tumours as described.^18^ BIA-ALCL seromas were obtained under ethics approval 17/ES/0047 awarded by the East of Scotland research ethics service. Samples are described in **Supplementary Table 1**. Diagnoses were confirmed by an independent histopathology review. Single-cell RNA sequencing was performed as previously described.^19^

### Flow sorting and 10x Genomics Chromium loading

Each vial of single cell suspension was thawed at 37°C. Cells were stained with CD45 and CD3 antibodies (both from Biolegend) as well as DAPI (1:100). Stained cells were then sorted using FACS into either of the following fractions: DAPI^-^ (viable cells), CD45^+^ CD3^+^ and CD45^+^ CD3^-^, with the goal of harvesting enough cells for 10x loading. A target of 2,000 to 10,000 cells was used to calculate the loading volume for the 10x Chromium, taken from the manufacturer’s protocol. Details for FACS and processed cell fractions can be seen in Supplementary Table 2. Each sorted fraction from every sample was loaded into one channel of the 10x Chromium chip before running on the Chromium Controller using 10x 5’ v2 kits.

### Spatial transcriptomics

For spatial transcriptomics, two paediatric cases of ALK-positive anaplastic large cell lymphoma (ALK+ALCL) were selected and processed using the Visium Spatial Gene Expression Kit for FFPE (10x Genomics, USA). Prior to analysis, RNA quality was assessed using the 4200 TapeStation System (Agilent Technologies, USA), with both samples showing DV200% values above 50%. Sample preparation was performed according to the manufacturer’s guidelines. Briefly, FFPE blocks with tissue dimensions of approximately 5x5 mm were prepared. Slides of 5-micron thickness were sectioned on a microtome and placed onto a Visium Gene Expression Slide with a capture area of 6.5x6.5 mm, containing 5000 total spots. Following deparaffinization, hematoxylin and eosin staining, and imaging were conducted. The subsequent steps included hybridization, ligation, release, and extension of the probes, followed by library construction. Sequencing was carried out on a NextSeq 550/500 platform (Illumina, USA). Sequencing depth was calculated based on the tissue section size on the capture area (>25000 read pairs/spot). For the first sample, the mean sequencing depth was 60000 read pairs/spot (2,130 median genes per spot), and for the second sample, it was 44000 read pairs/spot (2,370 median genes per spot).

### Drug Screens

Two commercially-available compound libraries were purchased from suppliers and used for high-throughput in vitro drug screens: the DiscoveryProbe™ Epigenetics Compound library (281 compounds; APExBIO, TX), and the FDA Approved Drug Screening Library (1430 compounds, Selleck Chemicals, TX). Biomek Software (v4.1) was used to program the liquid handling robot (Beckman-Coulter NXp Laboratory Automation Workstation). 96-well ’master’ drug plates (281 compounds from DiscoveryProbe™ Epigenetics Compound library, APExBIO, TX; and 1430 compounds from FDA Approved Drug Screening Library, Selleck Chemicals, TX) were diluted into 384-well ’mother’ plates on a per-quadrant basis, followed by two sequential dilutions to achieve a final per-drug concentration of 1μM. Cells were seeded at cell-line specific densities previously determined to maximise growth over the 48 hour incubation time of the screens. Adherent cell lines were seeded the day before addition of the diluted drug to allow time for attachment to the floor of the flask. CellTitre Blue (Promega) was added after 48 hours for a final 4 hours of incubation at 37°C/5%CO_2_. CellTitre Blue fluorescence was measured using a SpectraMax i3 (Molecular Devices; CA) plate reader using default settings for CellTitre Blue fluorescence. Fluorescence data were normalised per-plate according to positive (100% effect) and negative controls (0% effect).

### Immunohistochemistry for TIM3 and CD30

Immunohistochemistry was performed on formalin-fixed paraffin-embedded (FFPE) tissue from 9 independent ALK+ ALCL patients using standard protocols on a Discovery-XT staining device (Ventana). Following heat-mediated antigen retrieval, consecutive sections were stained for TIM3 (MAB23652, R&D Systems) and CD30 (M0751, Agilent Dako) both at a 1:50 dilution. Staining was developed under visual control. Images were captured with a Zeiss AxioImager Z1 and quantification performed with HistoQuest and StrataQuest (TissueGnostics GmbH, Vienna, Austria).

### Data deposition and analysis

The raw sequencing data generated in this study have been deposited in the European Genome-phenome Archive under accession number EGAD00001015703. Code generated is available at GitHub: https://github.com/ruoyan-li/T-cell-lymphoma. Supplementary Methods include descriptions of data analysis.

## Results

### A single cell dataset of T cell lymphoma entities

We sampled a range of TCLs from 18 individual patients from clinically realistic sources (tumour-involved lymph nodes (LN), malignant seromas and PDX including systemic/nodal ALK+ ALCL (n=3 PDX; NPM1-ALK translocated)^18^, BIA-ALCL (n=2 seromas), PTCL-NOS (n=5 nodal tumours), nTFHL-angioimmunobalastic (AI) (n=1 nodal tumour), nTFHL-NOS (n=1 nodal tumour), TLBL (n=5 nodal tumours) and one case of Sezary Syndrome (SS, involved nodal tumour of a sub-cutaneous malignancy)) (**Figure 1A; Supplementary Table 1**). Droplet-based 5’ scRNA-seq with TCR enrichment (10X Chromium platform) was performed for all live cells, or the viable CD45^+^CD3^-^ and CD45^+^CD3^+^ fractions following cell sorting by flow cytometry (**Figure 1A; Supplementary Table 2; Supplementary Figure 1A**).

**Figure 1.**
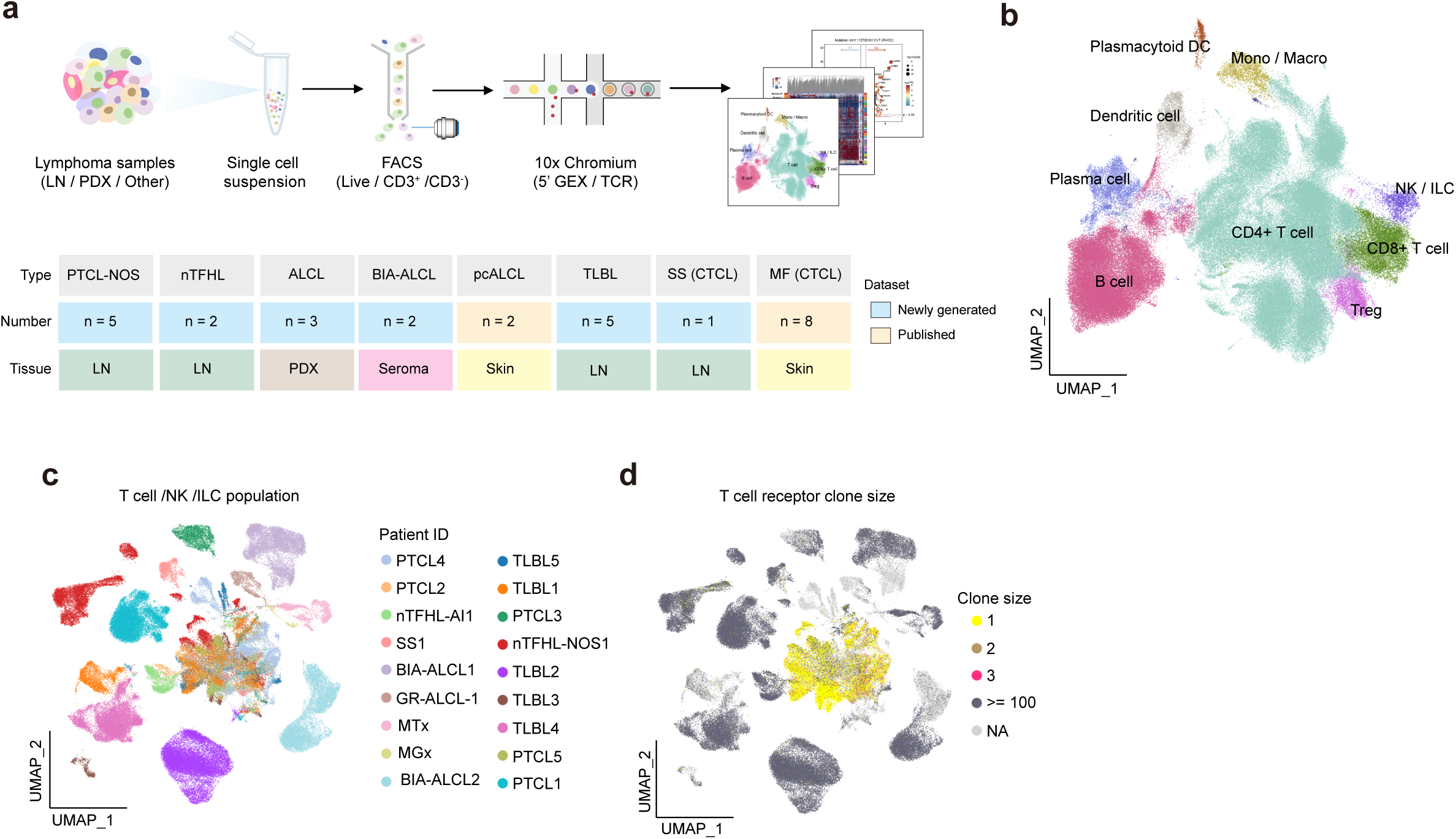
Overview of the dataset and T cell populations. (A) Schema of the experimental approach (top panel) and description of the tumour samples analysed, indicating which data were generated in this study and that taken from published datasets (bottom panel). Also summarised in Supplementary Table 1. (B) A UMAP showing the identities of the 180,332 cells captured across all samples. (C) A UMAP showing integrated data of the T cells and NK/ILCs from each of the patient samples representing a range of T cell lymphomas as indicated in (a). Each colour represents data from an individual patient tumour. PTCL4 and PTCL5 were excluded from this analysis due to a lack of identifiable malignant cells. (D) A UMAP showing clone size according to T cell receptor rearrangements across all captured T cells. PDX = Patient derived xenograft; LN = lymph node; PTCL-NOS = peripheral T cell lymphoma, not otherwise specified; nTFHL = nodal T follicular helper lymphoma; ALCL = anaplastic large cell lymphoma; BIA-ALCL = breast implant associated ALCL; TLBL = T lymphoblastic lymphoma; CTCL = cutaneous T cell lymphoma; MF = Mycosis fungoides; SS = Sezary Syndrome; TCR= T cell receptor; DC = dendritic cell; NK = Natural killer; ILC = innate lymphoid cell; GEX = gene expression.

Following data quality control steps and filtering, we captured 180,332 cells which were categorised into nine broad cell types based on the expression of canonical marker genes (**Figure 1B; Supplementary Figure 1B-D**). We added published scRNA-seq data of two other TCL sub-types for downstream comparisons: primary cutaneous ALCL (pcALCL, n=2) and mycosis fungoides (MF, n=8), the most common types of CTCL (**Figure 1A**). Whilst the majority of cells are of a T cell origin, we identified other cell types as indicated, which likely account for microenvironmental cellular infiltration (**Supplementary Table 2**).

To distinguish malignant from benign infiltrating T cells, we inferred large-scale chromosomal copy number variations (**Supplementary Figure 1E**). We found that malignant T cells from individual patients clustered separately, reflecting strong inter-donor heterogeneity (**Figure 1C; Supplementary Figure 1F**) as also observed for the disease entities (**Supplementary Figure 1G**). Two PTCL cases with no detectable malignant cells were excluded, but these data were used in subsequent microenvironmental analyses. Further projection of TCR clone size onto the UMAP revealed malignant T cells from each donor harboured a single clonally expanded TCR (**Figure 1D**), confirming they are malignant cells derived from clonal expansions of a single T cell.

### Integration of T cell development and TCR rearrangement status reveals insights into the cell of origin

We integrated malignant and benign T cells with foetal and adult thymic references using CellTypist to generate hypothesis-level origin assignments.^20,21^ We found the majority of TLBL cases aligned with double negative (DN) thymocytes, consistent with the established immature cell-of-origin and immunophenotype of this disease (**Figure 2A; Supplementary Table 1**).^22^ In comparison, ALK+ ALCL cases, although showing lower prediction probabilities, were found to align with double positive (DP) thymocytes for two of the three cases in keeping with a proposed thymic origin.^23–26^ In contrast, BIA-ALCL, like PTCL-NOS, presented as Tregs or other more differentiated cell types (**Figure 2A**). We integrated our malignant and benign T cell data with those from two published CTCL studies^19,27^, foetal T cell populations from a pan-foetal immune atlas^28^ and adult pan-body T cell data.^20^ For the most part, distinct TCL subtypes formed separate clusters, in accordance with the diagnoses previously made by histological interpretation (**Supplementary Figure 2A-C; Supplementary Table 1**).

**Figure 2.**
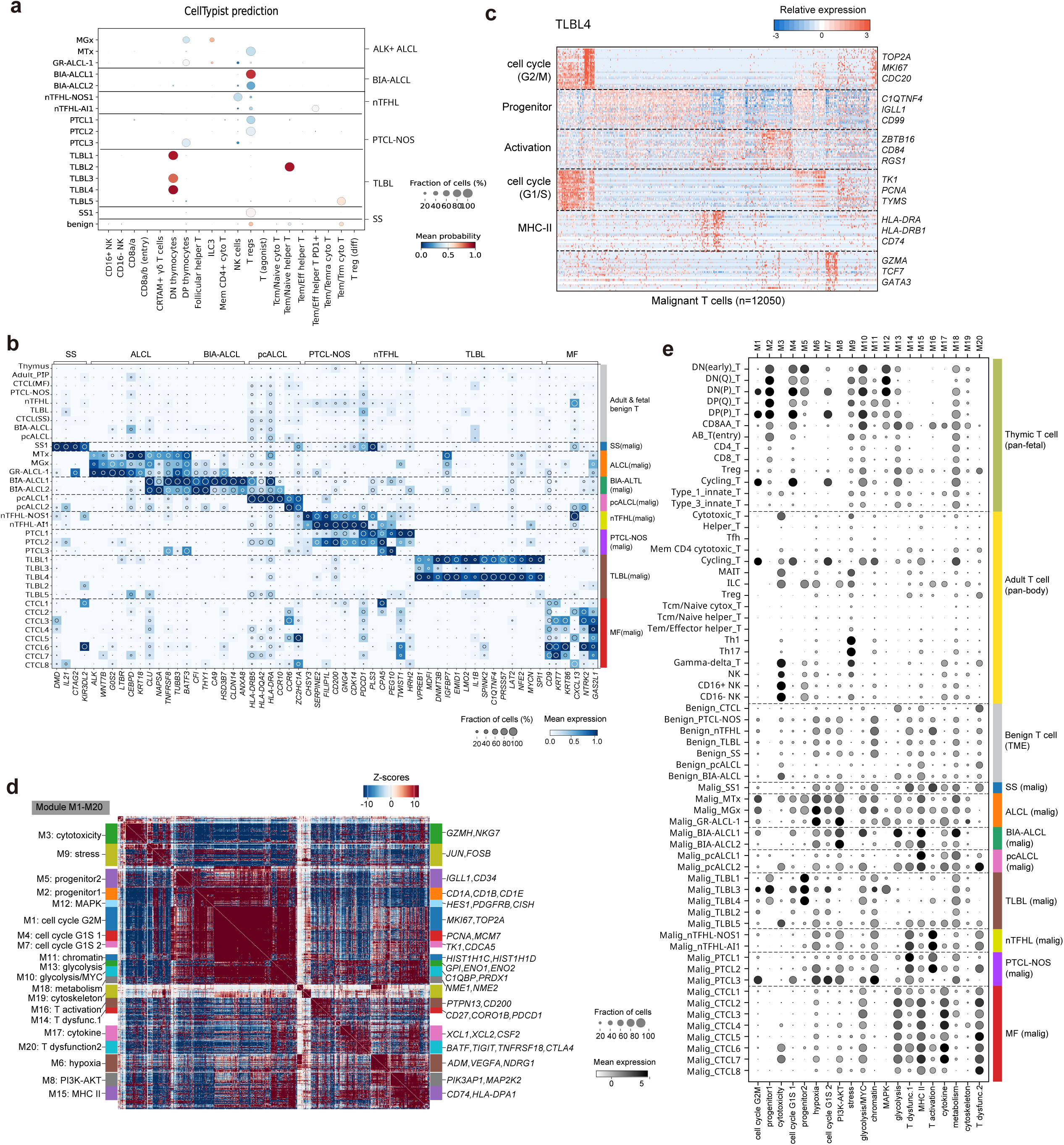
Comparison of disease entities and their transcriptional programmes reveals the potential cell of origin and descriptive genetic programmes. (A) CellTypist prediction of the cell of origin for each of the patient samples analysed, grouped according to histological entity. Only cell types for which query cells were assigned are shown. (B) A heat map of a curated subset of differentially expressed genes (DEGs) in malignant T cells according to disease classification. The depth of colour represents gene expression levels standardised between 0 (light blue) and 1 (dark blue). The inset circle indicates the percentage of cells expressing a given gene. (C) A heat map showing gene expression programmes and intra-tumour heterogeneity of malignant T cells in a representative patient (in this figure TLBL case 4). Programme annotations (left axis) and representative genes (right axis) are indicated as shown. The colour key indicates scaled expression levels (light blue, low expression to dark red, for high expression). (D) Gene modules identified in integrated benign and malignant T cell data using Hotspot. The heatmap comprises 1,000 highly variable genes with significant autocorrelation (FDR < 0.01), grouped into 20 gene modules on the basis of pairwise local correlation. (E) Dot plot showing gene module scores across samples (identified on the left axis by disease classification/benign cell subset shown on the right axis). The depth of shading of each dot indicates levels of gene module scores, ranging from white indicating no expression to dark grey depicting high expression. The dot size indicates the fraction of cells expressing a given gene module.

Given the association of some TCL with autoimmune diseases, we also aligned these datasets with an atlas of T cells present in patients with these conditions^29^ (**Supplementary Figure 2C**). The algorithm was unable to assign a cell of origin with high probability to any of the TCL, except most TLBL which aligned closely to SOX4+ T naive cells, in keeping with the stemness programme associated with SOX4 and immature T cell precursors (**Supplementary Figure 2D,E**).

To explore gene expression features of malignant T cells within each lymphoma subtype, we analysed DEGs across cancer types and between malignant and benign cells (**Figure 2B**).

DEGs distinguishing the TCL subtypes were as expected, whereby ALK+ ALCL uniquely expresses *ALK*, *TNFRFS8* (CD30) and *BATF3;* the latter two of which are shared with BIA-ALCL.^30^ In addition, BIA-ALCL, as previously reported, highly expresses the hypoxia gene *CA9.*^31^ Interestingly, pcALCL shared some features with BIA-ALCL, including expression of MHC class II genes *HLA-DRB5*, *HLA-DQA2* and *HLA-DRA,* perhaps suggesting some shared disease mechanisms.^32^ This signature was also seen in some CTCL cases, specifically MF, which was defined here by expression of *CD9*, *CXCL13*, *NTRK2* and *GAS2L1*. Interestingly, CD9 expression has previously been associated with tissue resident memory cells in autoimmunity, specifically Sjogren’s syndrome.^33^ Of the TLBL studied, two cases (TLBL1 and TLBL4) could be distinguished from the other TCL and benign T cells by expression of *MYCN*, *DNMT3B*, *LMO2* and *LAT2*. In particular, *LMO2* differentiates TLBL from benign lymphoproliferations as previously reported.^34^ Of the PTCL-NOS samples with malignant cells, PTCL1 and PTCL2 could be assigned to the PTCL-GATA3 subtype and PTCL3 was assigned to the PTCL-TBX21 subtype on the basis of gene expression.^35^

### Transcriptional programmes identify potential mechanisms of disease pathogenesis

We observed strong patterns of intra-tumour gene expression heterogeneity, exemplified by TLBL4 as shown in **Figure 2C**. Therefore, we investigated inter- and intra-tumour heterogeneity further through gene modules extracted utilising Hotspot.^36^ Through comparison with adult pan-body^20^ and foetal thymic T cells^28^, we aimed to capture gene modules upregulated in malignant T cells, representing tumour specific expression programmes; and modules shared between malignant and healthy mature/developing T cells reflecting cell types of origin or co-optation of developmental cell states in cancer. Twenty gene modules were annotated in total. (**Figure 2D; Supplementary Figure 2F; Supplementary Table 3**).

Taking the TCL cohort as a whole, we identified transcriptional programmes that were upregulated in malignant T cells relative to benign counterparts (**Figure 2D,E**). Module 16 was characterised by genes such as *PTPN13* and *CD200*, thus representing a gene programme related to T cell activation. We identified two potential T cell dysfunction-related modules, Modules 14 and 20, exemplified by genes such as *CD27* and *PDCD1*; and *TIGIT* and *CTLA4* respectively. The former showed high activity mainly in PTCL-NOS, while the latter was mainly active in MF (**Figure 2E**). Module 8 was characterised by genes like *PIK3AP1* and *MAP2K2*, and displayed high activity in PTCL-NOS and all entities of ALCL, in accordance with the known roles of the PI3 and MAP Kinase pathways in these malignancies.^5,37^

We detected five gene modules shared between foetal immature and malignant T cells, particularly in ALCL and TLBL (**Figure 2E**). Modules 1, 4 and 7 represent stages of the cell cycle, as reflected by genes such as *MIK67*, *PCNA* and *TK1* whereas Module 2 contains genes like *CD1A*, *CD1B* and *CD1E* and was shared between TLBL and thymic immature T cells (**Figure 2D,E**). Moreover, Module 5 consists of genes related to haematopoietic and T cell progenitors (e.g., *IGLL1* and *CD34*) and showed high activity in benign early DN T cells as well as the TLBL and ALCL cases. Therefore, this module may represent features of early immature T cells. Interestingly, high expression of Module 5 was also observed in three TLBL and two ALCL cases, reinforcing the high confidence prediction of a DN T cell origin for the former and DP T cell for the latter (**Figure 2A**).

### TLBL map to primitive thymic T cells yet produce transcripts associated with active TCR signalling

Projection of tumour cells onto the pan-foetal immune atlas T cell populations confirmed that three TLBL cases (TLBL1, TLBL3 and TLBL4) overlapped with primitive cell types (**Figure 3A,B**). To investigate the biology underlying this primitive phenotype further, we decided to focus our analysis on these three cases. We investigated expression of previously reported DN and progenitor T cell marker genes in these three cases and found high expression of *IGLL1*, *NFE2* and *HHEX* (**Supplementary Figure 3A**). Interestingly, malignant T cells from TLBL1 showed high expression of *CD34* (**Supplementary Figure 3A**), a well-known haematopoietic stem/progenitor cell marker. We subsequently incorporated TCR information using ‘Dandelion’ and found that malignant T cells in each of these three TLBLs shared a single non-productive TCR beta chain (**Supplementary Figure 3B,C**), indicating that they each stemmed from clonal expansion of a single T cell precursor. Overall, a lack of TCR V genes was widespread among these malignant T cells, consistent with a previous report that the majority of non-productive *TR* rearrangements are caused by absent V genes (**Supplementary Figure 3B**).^38^

**Figure 3.**
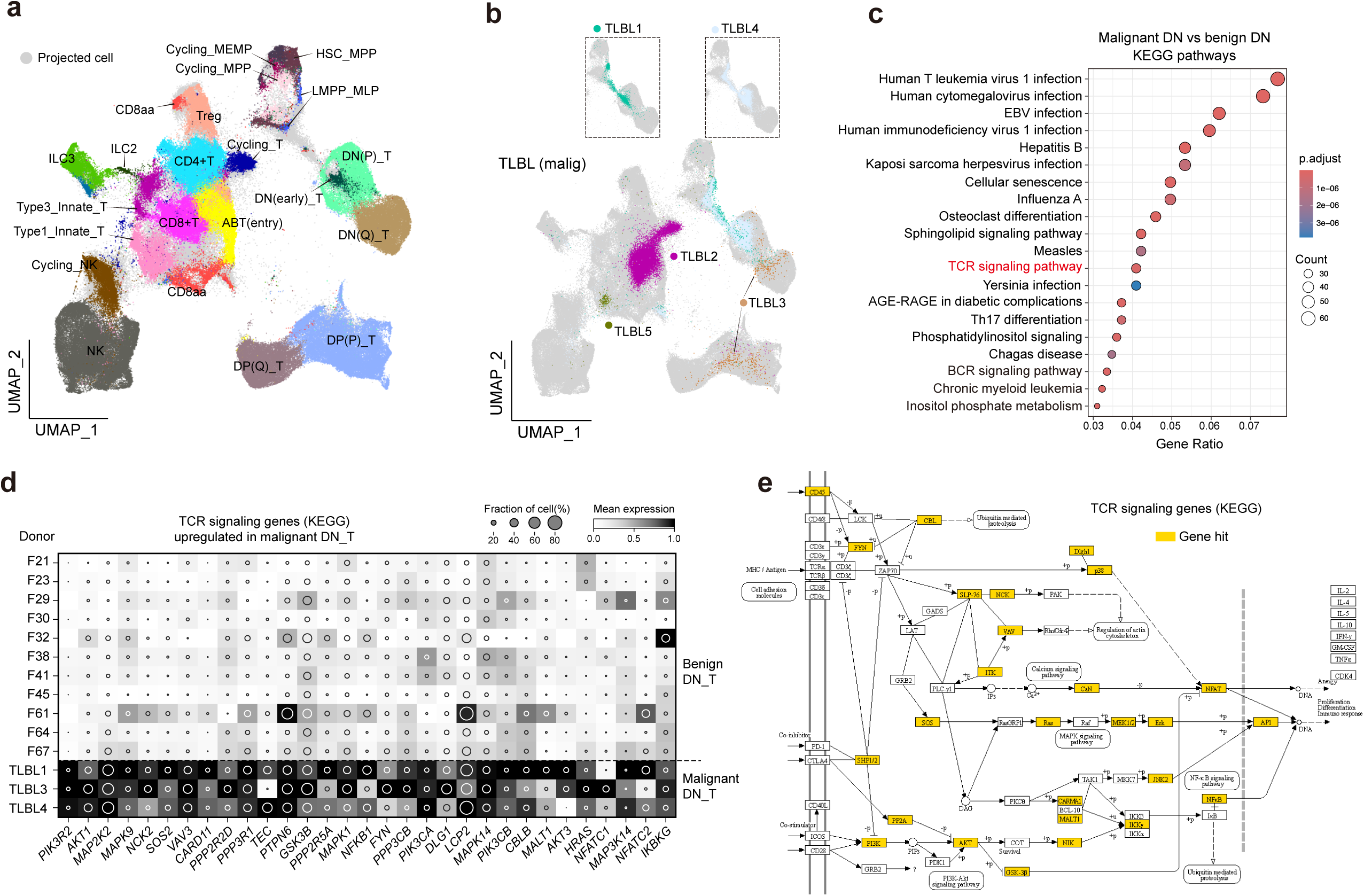
TLBL tumour cells map to thymic progenitor cells and upregulate expression of genes associated with T cell receptor signalling. (A) Projection of malignant T cells from our TLBL cases onto the UMAP of benign T cells from healthy thymus. (B) A UMAP showing the superimposition of malignant T cells from the TLBL cases onto benign T cells from healthy thymus. Individual tumours are highlighted. (C) KEGG pathway enrichment of upregulated genes in malignant double negative (DN) thymocytes (analysis restricted to TLBL1, TLBL3 and TLBL4) compared with the benign counterpart. The TCR signalling pathway is highlighted. (D) Heat map of TCR signalling pathway-related DEGs upregulated in malignant DN cells compared with the benign counterpart (analysis restricted to TLBL1, TLBL3 and TLBL4). The colour scale represents expression levels standardised from 0 (no expression) to 1 (expression). The inset circle indicates the percentage of cells expressing a given gene. (E) KEGG gene network of the TCR signalling pathway. Those DEGs upregulated in TLBL (malignant DN cells of TLBL1, TLBL3 and TLBL4) are highlighted in yellow.

We then compared malignant TLBL cells with their healthy DN counterparts^39^, focusing on the three cases with a predicted primitive origin. Employing pseudo-bulk DEG analysis, we observed viral infection and TCR signalling genes were upregulated in malignant DN T cells (**Figure 3C**). Due to the lack of productive TCRLJ rearrangement in these cases (**Supplementary Figure 3B**), upregulation of TCR signalling in malignant cells may reflect underlying oncogenic events rather than activation through TCR engagement. Indeed, distal pathways associated with an active TCR were present in the cells including PI3K, MAPK and NFKB (**Figure 3D,E**). Given functional TCRs are usually critical for T cell development, differentiation and activation; abnormal activation of the TCR signalling pathway implies oncogenic events in TCR-naked malignant thymocytes can mimic TCR signalling to empower their proliferation and activation.

### ALCL sub-types classified by pathways provides insights into mechanisms of disease pathogenesis

A defining feature of ALCL is expression of CD30 (*TNFRSF8*)^1^, which was detected in all cases, although subtype-specific genes were also seen (**Figure 2B**). To explore the latter further, we conducted KEGG enrichment and gene set enrichment analysis (**Supplementary Figure 4**). KEGG analysis revealed pathways associated with infectious processes across ALCL sub-types, suggestive of an activated cell phenotype (**Supplementary Figure 4A-C**). Of note, Th17 differentiation was a common hallmark; pathways associated with autoimmune conditions were identified for BIA-ALCL and pcALCL. Interestingly, comparison of BIA-ALCL to ALK+ ALCL showed the upregulation of hallmark gene sets associated with oxidative phosphorylation, hypoxia and allograft rejection (**Supplementary Figure 4D**).

Pathways associated with oestrogen and androgen signalling were enriched in pcALCL compared to BIA-ALCL, suggestive of a role for dysregulated oestrogen signalling in the former (**Supplementary Figure 4E**). The hallmark UV response was also a feature of pcALCL, consistent with skin localisation of this disease. Genes associated with the cell cycle (E2F targets) were upregulated in ALK+ ALCL compared to both BIA-ALCL and pcALCL, perhaps highlighting the more aggressive nature of systemic ALK+ ALCL compared to pcALCL (**Supplementary Figure 4F**).

### Intra-tumour heterogeneity: subclonal mutant proteins as potential drivers of plasticity and differentiation states in PTCL

To investigate intra-tumour phenotypic heterogeneity association with genotype, we detected somatic mutations from scRNA-seq data using Scomatic^40^ and reconstructed mutant clonal architectures. Overall, in most tumours, we observed that malignant T cells from different mutant clones mingled throughout UMAP space, indicating no overt phenotypic consequence brought by most mutations acquired (**Supplementary Figure 5**). However, we identified nine mutations in PTCL1, grouping 1,288 malignant T cells into four mutant clones and thus noticed malignant T cells from Clone 4 disproportionately distributed to one cluster (**Figure 4A,B**). This cluster showed an effector and cytotoxic-like phenotype, as reflected by high expression of *GZMA* and *GZMK,* as well as checkpoint receptors associated with exhaustion including *LAG3* and *TIGIT* (**Figure 4C,D**). Clone 4 was defined by an exonic nonsynonymous mutation in the gene *RHOC* (**Figure 4A**), a member of the Rho GTPase family that regulates the TCR and its co-stimulatory signalling pathway.^41^ The site of the variant itself is highly evolutionarily conserved (**Supplementary Figure 6A**) and the mutation scores in the top 0.1% most deleterious of all possible SNVs by CADD (**Supplementary Figure 6B**). The amino acid change (D59N) in the switch II region is predicted to destabilise an outer-sphere interaction with ionic magnesium, which normally participates in GTP hydrolysis (**Supplementary Figure 6C**). This could lead the molecular switch to a constitutive “on” state by blocking GTP hydrolysis. Thus, we identify a RHOC-mutant subclone potentially associated with cytotoxicity/exhaustion; causality and prevalence across PTCL-NOS remains to be validated.

**Figure 4.**
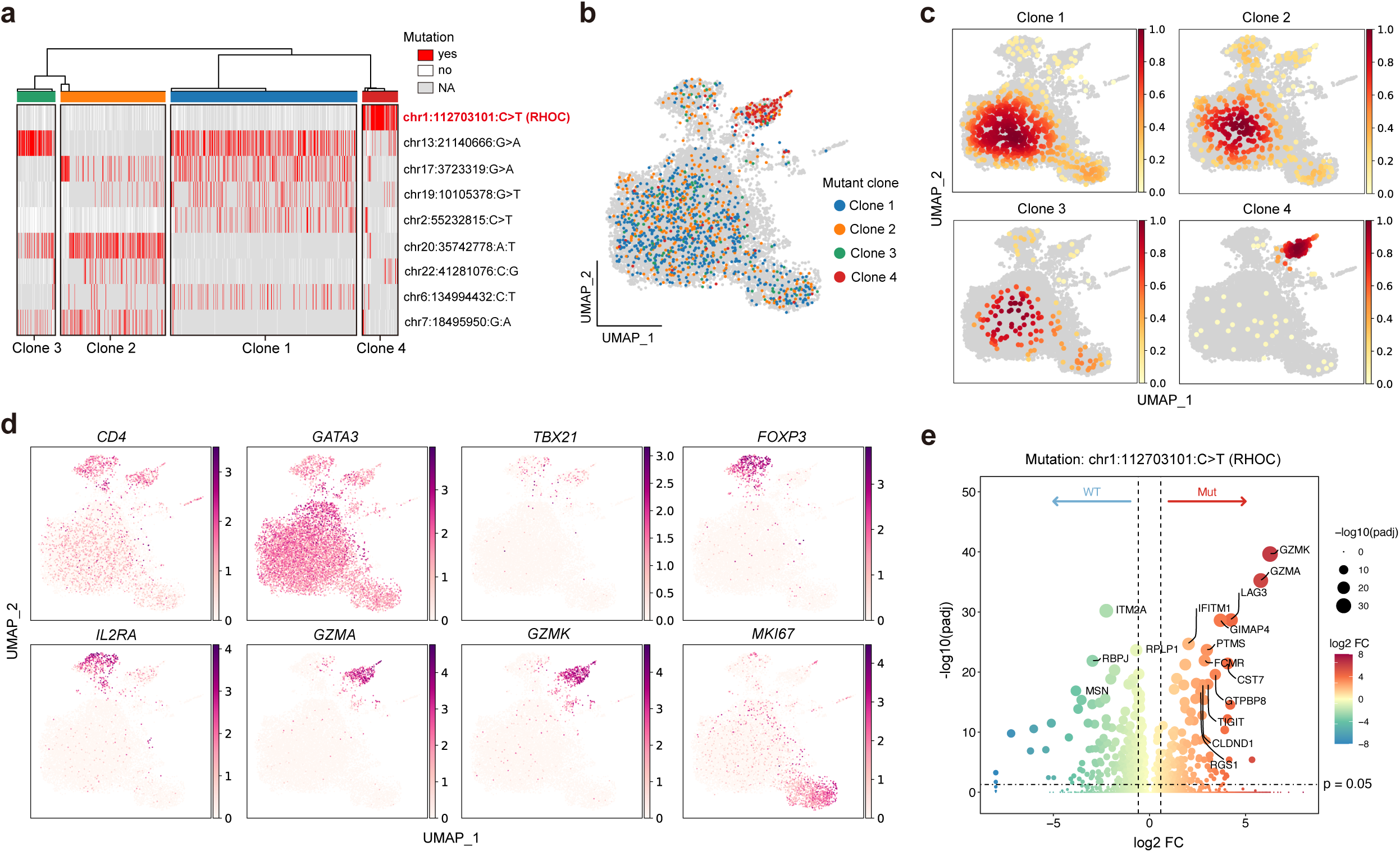
Intra-tumour genotypic and phenotypic heterogeneity in PTCL reveals a cytotoxic T cell sub-clone expressing a potential novel oncogenic *RHOC* mutation. (A) A heat map of somatic mutations called from scRNA-seq data of a representative tumour (PTCL1). Malignant T cells are clustered into four clones. (B) A UMAP of malignant T cells from PTCL1 coloured by genotypic clones. Grey shading represents unassigned clonal information. (C) Density plots showing the distribution of malignant T cells from different genotypic clones in the expression feature space. (D) UMAPs of DEGs in different cell clusters of PTCL1. (E) Volcano plot showing DEGs between malignant T cells with mutation in the *RHOC* gene (the defining mutation of clone 4) and other cells.

### Spatial analysis of the tumour microenvironment unveils TIM3 as an immunotherapeutic target for TCL

To understand TME changes in TCL, we integrated our data with that from healthy lymph nodes from a pan-body atlas.^42^ CellTypist annotated ten broad cell types and 29 cell subtypes, of which effector memory/resident memory cytotoxic T cells, Tregs, proliferative germinal centre B cells and migratory DCs had increased abundance in diseased lymph nodes (**Supplementary Figures 7A,B**). This is consistent with an inflammatory response to the tumour and concurrent down-regulation of the immune response enacted by Tregs (**Supplementary Figures 7C,D**).

As the ALK+ ALCL cases subjected to scRNAseq were derived from PDX lacking the TME, we also analysed two independent primary tumour sections using spatial transcriptomics (**Supplementary Figures 8, 9**). Briefly, we integrated scRNA-seq from ALCL PDX tumours with two healthy lymph node samples^42,43^ (**Figure 5A,B)** and used cell2location to predict cell type abundances captured by 10X Visium (**Supplementary Figures 10A, SB**). The samples showed extensive invasion by neoplastic T cells, replacing large areas of lymphoid tissues with scattered residual germinal centers (**Figure 5C,D**). Of note, TIM3^+^ infiltrating T cells were predicted to be enriched in areas surrounding tumour cells (**Figure 5C,E**). A known co-regulator of immunosuppression, we found TIM3 expression on 30-70% of infiltrating T cells of a further 9 independent cases of ALK+ ALCL by immunohistochemistry (**Figure 5F,G**). These data suggest that TIM3 inhibitors might be a viable therapeutic approach for ALK+ ALCL. We then assessed our scRNA-seq TCL dataset for expression levels of TIM3 and other immune checkpoint proteins on infiltrating T cells, and their ligands on tumour cells (**Supplementary Figures 11A,B**). *LAG3* was particularly prevalent on benign T cells in PTCL, as was *TIGIT*, the latter of which was also a feature of TLBL although to a lesser extent (**Supplementary Figure 11A**). These data suggest that immunotherapeutic approaches should be explored further for other TCL entities.

**Figure 5.**
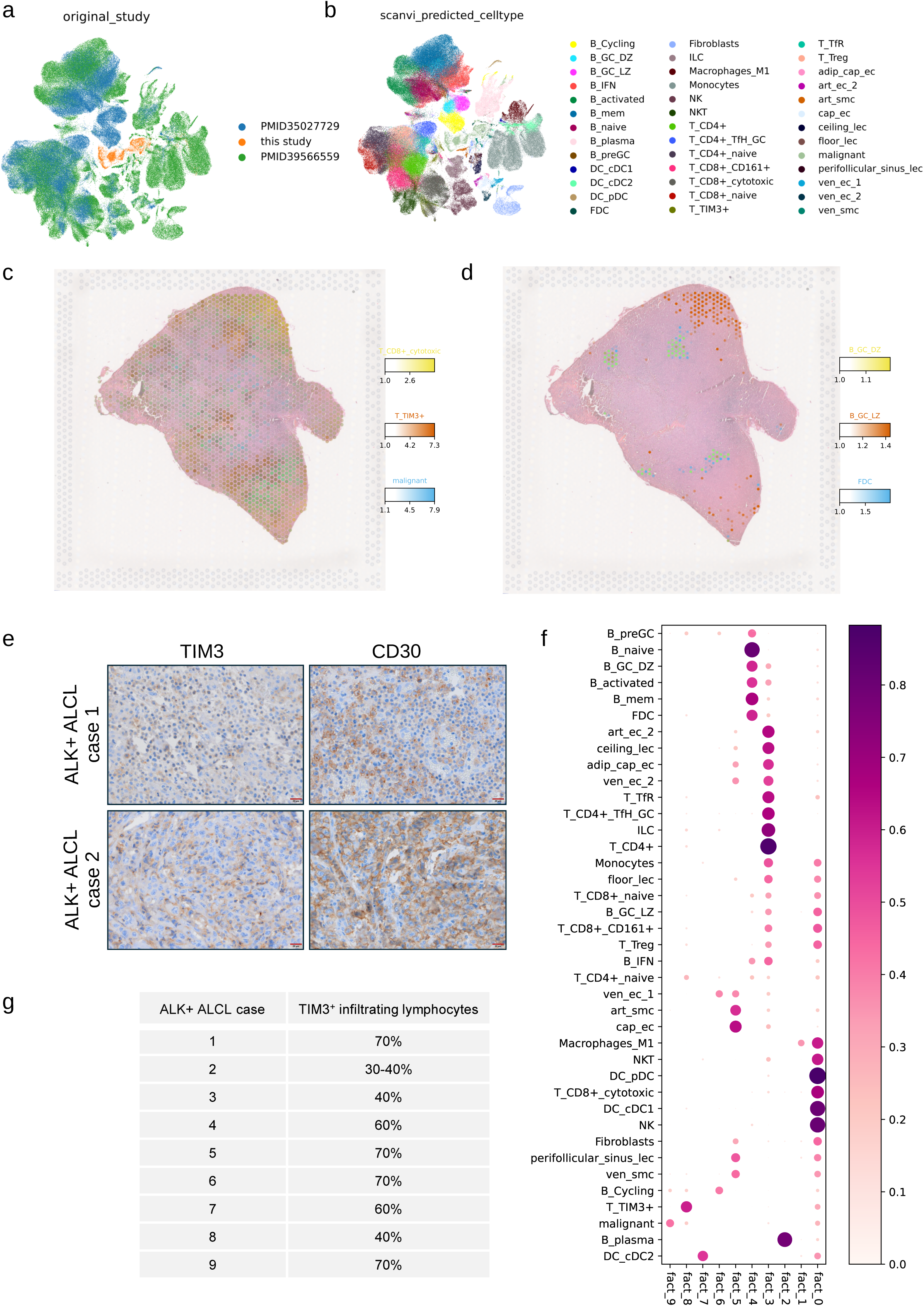
A suppressive immune microenvironment is present in ALK+ ALCL exemplified by TIM3 expression on infiltrating T cells. (A) A UMAP showing the integration of tumour and benign cells of the TME, and healthy lymph node data from two public studies (PMID35027729, PMID39566559). Cells are coloured according to the data source, and scANVI-predicted cell types are shown in (B), (C) and (D). Images of an ALK+ ALCL case colored by Cell2location-predicted cell type abundances in each Visium spot. (C) Predictions for the locations of malignant cells, TIM3+ T cells (T_TIM3+) and cytotoxic T cells (T_CD8+_cytotoxic), (D) Predictions for the locations of dark zone B cells (B_GC_DZ), light zone B cells (B_GC_LZ) and follicular dendritic cells (FDC). (E) A dotplot showing non-negative matrix factorisation (NMF) factors, where each factor is a group of co-localized cell types. In the dotplot, the size and color density represent the loading of each cell type in each factor. (F) Staining of TIM3 and CD30 expression on two representative ALK+ ALCL cases (G) Summary of the % TIM3+ infiltrating T lymphocytes in each of the 9 ALK+ ALCL cases analysed.

### Benchmarking therapeutic target prediction using patient-derived models reveals a potential role for SYK as a therapeutic target in TCL

We further investigated potential therapeutic targets active in malignant T cells using drug2cell analysis, which predicts therapeutic targets leveraging the ChEMBL database^44^ (**Figure 6A**). To benchmark for predictions against actual biological activity, we conducted an *ex vivo* drug screen of ALCL PDX using two drug libraries collectively containing over 1,700 drugs. Comparison of the drug screen data to the drug2cell analysis for ALCL demonstrated significant correlation (p < 2×10^-16^, p = 3.8×10^-9^, p = 3.1×10^-6^ for MGx, MTx, and GR-ALCL-1 ALCL PDX respectively), suggesting that drug2cell scores can be predictive of *in vitro* drug sensitivity (**Figure 6B**). For example, drugs with known therapeutic efficacy in ALK+ ALCL, such as crizotinib and ceritinib, were predicted *in silico*, as were cytotoxic therapeutic agents such as vincristine and vinblastine. The latter agents were predicted by the drug2cell analysis to have efficacy in the treatment of the other TCLs as expected (**Figure 6C**). Hence, the predictions produced for PTCL, for which few good pre-clinical lab models exist, may have biological significance, which is particularly pertinent given the paucity of efficacious therapeutic options. In particular, this analysis revealed potential sensitivity for inhibitors of mTOR, XPO1, CDK, EGFR and HDACs in the treatment of PTCL (**Figure 6C**).

**Figure 6.**
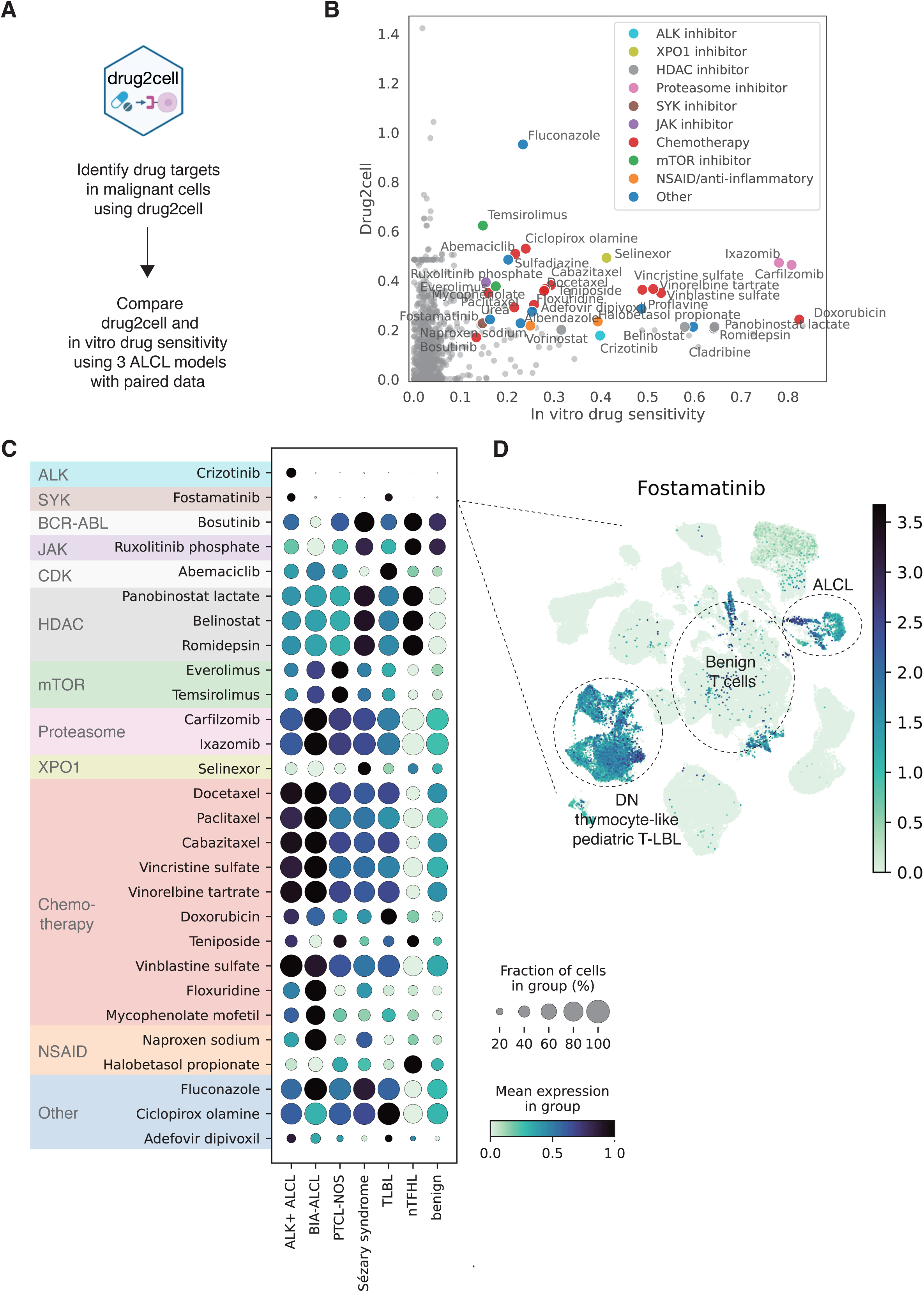
Prediction of therapeutic targets reveals a potential role for SYK and other targeted inhibitors in the treatment of ALCL and T-LBL. (A) The workflow for the integrated analysis of drug2cell drug target prediction and in vitro drug sensitivity data. (B) A scatter plot comparing average drug2cell scores, and average in vitro drug efficacy for the 3 ALCL PDX models analysed. Drugs showing both a drug2cell score and an in vitro drug sensitivity of over 0.25 are coloured. (C) A dot plot of drug2cell scores across cancer types and benign T cells for the selected drugs. (D) A UMAP projection of all scRNAseq TCL patient data coloured by fostamatinib prediction scores.

Intriguingly, the SYK inhibitor fostamatinib was identified by the drug2cell analysis as a potential therapeutic approach for ALCL and TLBL (**Figure 6C**). Given our finding of increased TCR signalling in these cases and the association of SYK with the pre-T cell receptor, SYK inhibition may represent a strategy to specifically target malignant cells with an early thymic phenotype. This was confirmed by the *ex vivo* drug screen for ALK+ ALCL whereby the SYK inhibitor fostamatinib led to a decrease in cell survival (**Figure 6C**). We validated this finding by exposing ALCL PDX cells *ex vivo* to fostamatinib (**Supplementary Figure 12**).

## Discussion

TCL are a heterogeneous group of malignancies considered to have derived from immature or mature T cells, although a thorough evaluation of disease mechanisms of translational relevance has been hampered by the heterogeneity of these malignancies and their relative rarity. To improve this understanding, we developed an exploratory single cell dataset of TCL which provides insights into disease pathogenesis and highlights therapeutic targets for future validation.

We first harnessed this dataset to explore disease mechanisms. Transcripts for *TR*-β rearrangements could be detected for all cases except TLBL, which matched subsequent CellTypist prediction and previous reports of a thymic origin for this disease.^45^ Although CellTypist predicted a thymic DP T cell of origin for ALK+ ALCL, this was not reflected in a lack of *TR* rearrangements, suggesting either a thymic origin with peripheral transformation, or NPM1-ALK-induced upregulation of an immature gene expression programme in mature

T cells as previously proposed.^25,46,47^ We also corroborate the previously reported capacity for NPM1-ALK to mimic proximal TCR signalling^37^ by activating distal pathways, describing here expression of a PI3K-AKT module in ALCL. Furthermore, it is considered that PTCL is sustained by active-TCR signalling, either through activating mutations in proximal TCR signalling proteins or other external factors engaging the TCR.^48–50^ Indeed, we see upregulation of a T-cell activation module in PTCL-NOS and nTFHL, suggestive of chronic active TCR-induced signalling playing a pathogenic role in these diseases. In contrast, whilst KEGG analysis indicated up-regulation of the TCR signaling pathway for our TLBL cases (**Figure 3C**), it has been reported for TALL that stimulation of cells through the CD3/TCR complex induces cell death in a manner akin to negative selection.^51^

Of interest, KEGG analysis of BIA-ALCL samples showed upregulation of a number of pathways associated with infection including viral, bacterial and parasitic as well as autoimmunity and allergy, all previously proposed as disease mechanisms.^52^ Prior reports have postulated cells of origin ranging from Th1/2 to Th17/Treg.^52,53^ Our data do not resolve these conflicting reports, as both Th17 and Th1/2 differentiation pathways show enrichment, although CellTypist predicted a Treg phenotype for BIA-ALCL (**Figure 2A**). In addition, we were able to sub-classify BIA-ALCL amongst other ALCL based on expression of the hypoxia factor CA9, which differentiated it from pcALCL as did up-regulation of gene sets associated with androgen/oestrogen signalling seen in the latter and has been previously reported for SS and MF in the context of ERLJ.^54^ All forms of ALCL shared expression of CD30 and BATF3 in keeping with other data^1,30,53^ and while our ALCL samples were derived from primary patient material as well as early passage PDX, we consider that cross-comparisons are still valid given the PDX represent close to 100% tumour cells as do seromas obtained from patients with BIA-ALCL.

Interestingly, we were able to detect a novel sub-clonal mutation in one PTCL-NOS case that would likely not have been detected by whole genome sequencing in routine clinical practice. The *RHOC* mutation observed is in a conserved region of the protein and bears similarities to mutations in the RHOA switch II region, such as Q63L, which have been experimentally demonstrated to lead to impaired hydrolysis and constitutive activation.^55–57^ *RHOC* itself is hyperactivated in a variety of other cancers albeit largely by overexpression,^58^ so we hypothesize that the *RHOC* mutation in this case might be a novel oncogenic driver in PTCL.

Our data also point towards a suppressive inflammatory microenvironment in TCL that could be exploited therapeutically. In particular, we compared Tregs in diseased and healthy lymph nodes and found higher expression of *PDCD1* (PD1) on Tregs in some cases; a finding previously associated with enhanced immunosuppression on PD1 blockade, particularly when PD1 expression levels are low on effector T cells^59^. This may present an important clinical consideration counteracting the use of nivolumab, indeed limited responses with hyper progressions have been reported for PTCL treated with nivolumab.^60^ Alternative immune checkpoint inhibitors for the treatment of PTCL have been less forthcoming due to the aforementioned issues of disease rarity and good quality samples for analysis.^61,62^ However, our study has highlighted LAG3 and TIGIT as potential therapeutic targets for the treatment of PTCL-NOS that should be further validated.

Given the scarcity of targeted therapeutic approaches available for most TCL (with the exception of ALCL), we explored our data using drug2cell and validated this approach through a high throughput *ex vivo* drug screen employing PDX-derived ALCL cells. This approach validated SYK as a therapeutic target in TLBL and ALCL as has previously been suggested in the case of ALCL and PTCL^63–65^ These data were validated here using *ex vivo* ALK+ ALCL PDX treated with the SYK inhibitor fostamatinib, but we could not corroborate the previously published expression of *SYK* in other PTCL in our cases suggesting this to be a sub-type specific therapeutic vulnerability, or might be detected if a larger number of cases are assessed in future.^65–67^

The correlation between drug2cell predictions and drug screen results suggest that targets predicted by the drug2cell methodology might represent viable therapeutic targets for follow-up in the absence of good pre-clinical PTCL models to test them. Indeed, drug2cell predicted the HDAC inhibitors romidepsin and belinostat as being efficacious in the treatment of PTCL in keeping with their approval for the treatment of relapsed/refractory PTCL.^68,69^ Furthermore, the mTOR inhibitor everolimus, also predicted by our analysis, has also been assessed for the treatment of PTCL^70^ as has the JAK1/2 inhibitor ruxolitinib.^71,72^ Finally, given the poor prognosis for these patients, our data suggest that other targeted drugs identified by us, such as XPO1 inhibitors^73^, perhaps combined with immune checkpoint inhibitors as discussed above, might be worthy of further investigation.

Due to the rarity and extensive heterogeneity of TCL, future studies of more cases, particularly in disease areas underrepresented in this dataset, will be required to draw broad general conclusions.

## Authorship

Conceptualization: S.D.T., S.A.T. and C.S. Methodology: R.L., J.D.M., M.S., Formal analysis: R.L., J.D.M., E.R.J., C.V.A., O.D., S.L., D.A., E.S., A.M.T., E.V., M.M.T., M.Q.D. Investigation: R.L., J.D.M., E.R.J., C.V.A., O.D., S.L., D.A., E.S., A.M.T., E.V., M.M.T. Resources: S.T., S.D.T., L.K., M.S., O.M., R.J., P.T., M.Q.D. Writing: S.D.T, R.L., J.D.M., E.J., C.V.A., Supervision: S.D.T. and S.A.T. Funding acquisition: S.D.T. and S.A.T.

## Conflict-of-interest disclosure

S.A.T. is a scientific advisory board member of ForeSite Labs, OMass Therapeutics, Qiagen, Element Biosciences, Xaira Therapeutics and Bioptimus, co-founder and equity holder of TransitionBio and EnsoCell Therapeutics, a non-executive director of 10x Genomics and a part-time employee of GlaxoSmithKline. All other authors declare no competing financial interests.

## Notes

https://github.com/ruoyan-li/T-cell-lymphoma

https://ega-archive.org/studies/EGAS00001005750

### Funding

- Cancer Research UK: CTRQQR-2021\10001, C9685/A25117
- Medical Research Council: MR N013433-1
- Children's Cancer and Leukaemia Group: CCLGA 2021 12
- European Union: 101072735EU
- NIHR Cambridge Biomedical Research Centre: BRC-1215-20014
- Wellcome Trust: WT206194

