## Supplementary figures for "Single-Cell Profiling Reveals Developmental Trajectories and identifies SYK and TIM3 as Targets in some T Cell Lymphomas"

**Supplementary Methods**

**Library preparation and sequencing**

Gene expression libraries were generated from the resulting cDNA after clean up following 10x Genomics protocols. TCR cDNA was enriched from each CD45^+^ and DAPI^-^ cell fraction and subsequent libraries were made from enriched TCR cDNA according to the manufacturer’s protocol. All libraries were sequenced using an Illumina NovaSeq.

**Single cell RNA-seq data processing, quality control and doublet removal**

Gene expression data from 10x Genomics droplet-based sequencing were aligned to the GRCh38 reference genome (GRCh38-2020-A) using the 10x software package CellRanger (version 6.1.2). The V(D)J data were processed using CellRanger (version 6.1.2) and the reference vdj_GRCh38_alts_ensembl-5.0.0 with default parameter settings. After alignment, we ran the SoupX algorithm with default parameters for 10x each run and applied adjustCounts() function to remove ambient mRNA from the count matrix. The outputs from SoupX were further read using the read_10x_mtx() function in Scanpy (version 1.8.1). We then concatenated data objects from different 10x lanes using the concatenate() function in anndata (version 0.8.0). To detect and remove doublets from the data, we applied Scrublet (version 0.2.3) to the data from each 10x lane to obtain per-cell scrublet scores and used a doublet exclusion threshold of median plus three median absolute deviations of the doublet score. Cells with greater than 40% mitochondrial gene expression or expression of fewer than 200 detected genes were excluded from downstream analysis. Genes that were expressed in fewer than 3 cells were also removed.

**Data normalisation, embedding, visualisation, clustering and integration**

We performed data normalisation to correct for cell-to-cell variation using the normalize_per_cell() function in Scanpy (version 1.8.1). Normalised data were then transformed using the log1p function in Scanpy to alleviate skewness of data and mean-variance relationship. Expression values of each gene were then scaled and centred using the scale() function in Scanpy. Highly variable genes (HVGs) were detected using the highly_variable_genes() function in Scanpy with minimum cut-off values 0.0125 and 0.5 for expression and dispersion. We then calculated principal components (PCs) using the pca() function in Scanpy. For batch correction, we used the harmonypy package to correct donor-to-donor variation with the theta value being set to 3. The neighbours() function was used to calculate the neighbourhood graph. UMAP embedding was calculated using the umap() function in Scanpy. The neighbourhood graph was then clustered using the leiden() function in Scanpy. Broad cell types were annotated based on expression of canonical marker genes.

**Inferring copy number variations based on scRNA-seq data**

To distinguish malignant T cells and non-malignant cells, we inferred large-scale chromosomal copy number variations of single cells based on scRNA-seq data using the tool InferCNV (https://github.com/broadinstitute/inferCNV) with default parameters. Briefly, InferCNV first orders genes according to their genomic positions (ﬁrst from chromosome 1 to X and then by gene start position) and then uses a previously described sliding-average strategy to normalise gene expression levels in genomic windows with a fixed length. Multiple putative non-malignant cells are chosen as the reference to further de-noise the CNV result.

**Cell type annotation with CellTypist**

We used CellTypist, an established method for automated reference-based cell type annotation to explore the proximity of malignant cells to physiological immune cell types.^1^ Briefly, CellTypist trains logistic regression classifiers based on annotated reference datasets and classifies query cells as the highest probability cell type present in the reference. To generate a custom CellTypist reference, we leveraged data of T cells from thymus and blood ^1^, and randomly sub-sampled to 0.3 of the total number of cells. We trained the model using the ‘celltypist.train’ function with a feature selection step. Prior to classification with CellTypist, we filtered out cells with mitochondrial read expression >=10% of total cell read count.

**TCR data analysis using Dandelion**

We applied Dandelion (version 0.3.2), which delivers improved V(D)J contig annotation and the identification of non-productive and partially spliced contigs, to analyse the TCR data in our study. Briefly, we first pre-processed the VDJ data using the Singularity container provided by Dandelion, which performs VDJ gene annotation with igblastn and parses to AIRR format. We then read in the files with the full contig space (all_contig_iblast_db-all.tsv) and concatenated them using the concat() function in Dandelion. Gene expression and VDJ objects were further linked using the check_contigs() function with ‘productive_only = False’ and ‘library_type = tr-ab’. For TCR clonotype calling, we ran the find_clones() function with ‘identity = 1’ and ‘key = junction’. After this, we quantified TCR clonal expression using the clone_size() function with ‘max_size = 100’. Finally, we copied the information from the Dandelion object to the Scanpy gene expression object using the transfer() function.

**Differentially expressed gene analysis on malignant and benign double negative T cells using a pseudo-bulk strategy**

We applied a pseudo-bulk strategy to the analysis of differentially expressed genes (DEGs) between malignant and benign double negative (DN) T cells. Briefly, we first aggregated raw counts of each gene by donor and thus used donors rather than cells as biological replicates in the DEG analysis. We then leveraged the edgeR R package for the DEG analysis. We filtered genes by expression levels using the filterByExpr() function in edgeR with ‘min.count’ and ‘min.total.count’ being set to 50 and 100 respectively. We designed the model matrix using the model.matrix() function and only included one variable, namely groups (malignant and benign DN T cells). We then fit genewise negative binomial generalised linear Models with quasi-likelihood tests using the glmQLFit() and glmQLFTest() functions in edgeR. The generated DEGs were further used for pathway enrichment analysis using the clusterProfiler R package. In particular, we focused on gene enrichment in Gene Ontology BP (Biological Process) using the enrichGO() function and KEGG pathways using the enrichKEGG() function.

**Investigation of gene modules in malignant T cells, and benign adult and foetal T cells**

To identify gene modules that are active in malignant T cells from different lymphomas and compared to benign adult and foetal T cells, we first merged the data of T cell populations (both malignant and benign T cells) in our study, those from two previously published cutaneous T cell lymphoma studies ^2,3^ and foetal T cell populations from a pan-foetal immune atlas^4^. We integrated these datasets using the scVI module within scvi-tools (version .19.0). Specifically, to develop a low-dimensional representation of the expression dataset, scVI was run on the 2000 highly variable genes with n_layers. We then applied HotSpot to identify gene modules in the integrated data object.^5^ We used HotSpot for this analysis mainly because it enables the identification of informative gene modules, with high locally autocorrelated genes, by examining the pairwise local correlation between genes on cell neighbourhoods in the nearest neighbour graph. This method has been reported as suitable for scRNA-seq data where gene drop-out in individual cells is common. The latent space from scVI was then used as input to Hotspot and modules were created from the top 1000 genes based on local autocorrelation (FDR < 0.01). For module creation, an fdr_threshold of 0.05 was used along with a min_cluster_genes setting of 20. We annotated these modules based on the 3CA database^6^ and gene set enrichment analysis.

We then identified a subset of significantly auto-correlated features, with respect to the scVI latent space, by running HotSpot using the depth-adjusted negative binomial (danb) observation model and 30 neighbours. The danb model comprises the background null distribution against which expression counts are normalised, and is used to avoid flagging genes as significant due to local autocorrelation in the library size of cells.

**Somatic mutation calling and mutant clone clustering**

We called somatic mutations from our scRNA-seq data using Scomatic, a method for identifying somatic mutations in high-throughput single-cell genomics and transcriptomics datasets.^7^ Briefly, we implemented Scomatic into a Nextflow pipeline to accommodate data from multiple 10x runs from the same patient. To reconstruct the clonal architecture of specific tumours, we first ran Scomatic on scRNA-seq data from each patient and then genotyped for detected mutations across cell types. We then used a matrix to record the status (wide type, mutated and NA) of each mutation detected in malignant T cells from each tumour. The matrix was further used to perform unsupervised hierarchical clustering. In the clustering analysis, we only included cases in which we could detect the presence of mutations in at least 50 malignant T cells and that could be genotyped in at least 2% of all malignant T cells. CADD, AlphaMissense and MetaRNN scores were computed for RHOC D59N and RHOA Q63L mutations using publicly available web tools (CADD: [cadd.gs.washington.edu/snv](https://cadd.gs.washington.edu/snv), MetaRNN: [liulab.science/metarnn](http://www.liulab.science/metarnn.html), AlphaMissense: alphamissense.hegelab.org).

**Spatial transcriptomics data analysis**

Primary data analysis (RNA-seq output and bright field microscope images to align reads, generate feature-spot matrices, perform clustering and gene expression analysis) and visualisation were performed using 10x Genomics Space Ranger 2.0.0 and Loupe Browser 6.4.1. Advanced bioinformatics analysis involved cell-type deconvolution with cell2location and co-localization analysis. For running cell2location, we integrated three different scRNA-seq datasets to build the reference, including a vascular atlas^8^ subsetted to relevant cell types in lymph nodes, the cell2location lymph node reference^9^ and the scRNA-seq data of the three ALCL PDX analysed here. scVI and subsequently scANVI were employed to harmonize cell type labels for the integrated reference dataset. After building the reference, a cell2location spatial mapping step was conducted with the following hyper-parameter choices: n_cells_per_location being 30 and detection_alpha being 200. Cell type colocalization was assessed using Non-Negative Matrix Factorization (NMF) with the cell2location package using the predicted cell abundance.

**Determination of EC50 values**

Fostamatinib Disodium [CAS 1025687-58-4] powder was reconstituted in PBS, 2% DMSO at 10mM and serially diluted into a 96-well stock plate. Drug stock (5μL) was added to 95μL of cell suspension using a NXp Laboratory Automation Workstation to achieve final drug concentrations ranging between 1nM and 100μM. Following 48 hours of incubation with the drug, 20μL of CellTitre Blue (Promega) was added and cells incubated for an hour at 37℃/5%CO_2_. CellTitre Blue fluorescence was acquired by the same method as for drug screening and normalised in the same way. Data was analysed using GraphPad Prism 10.2.1 with a two-parameter log-logistic function (log(inhibitor) vs normalized response).

**Drug2cell analysis**

Drug2cell^11^ was run on the combined object containing both malignant and benign T cells. For ALK inhibitors, of specific interest in ALCL, the target gene list was revised to exclude fusion partners NPM1 and EML4 to focus the score on the actual kinase targets (ALK) for more accurate results. Single-cell level drug2cell scores were pseudobulked for each patient sample using decoupler^12^. To identify drug targets supported both by drug2cell and in vitro data, drugs were subsetted to ones with both a drug2cell score and an in vitro drug sensitivity score >0.25 in at least one of the three PDX models, with both scRNA-seq and in vitro drug sensitivity data available. The resulting drugs were examined grouped by disease and patient to identify drugs with a disease or patient subset-specific target expression pattern.

**Supplementary Figures**

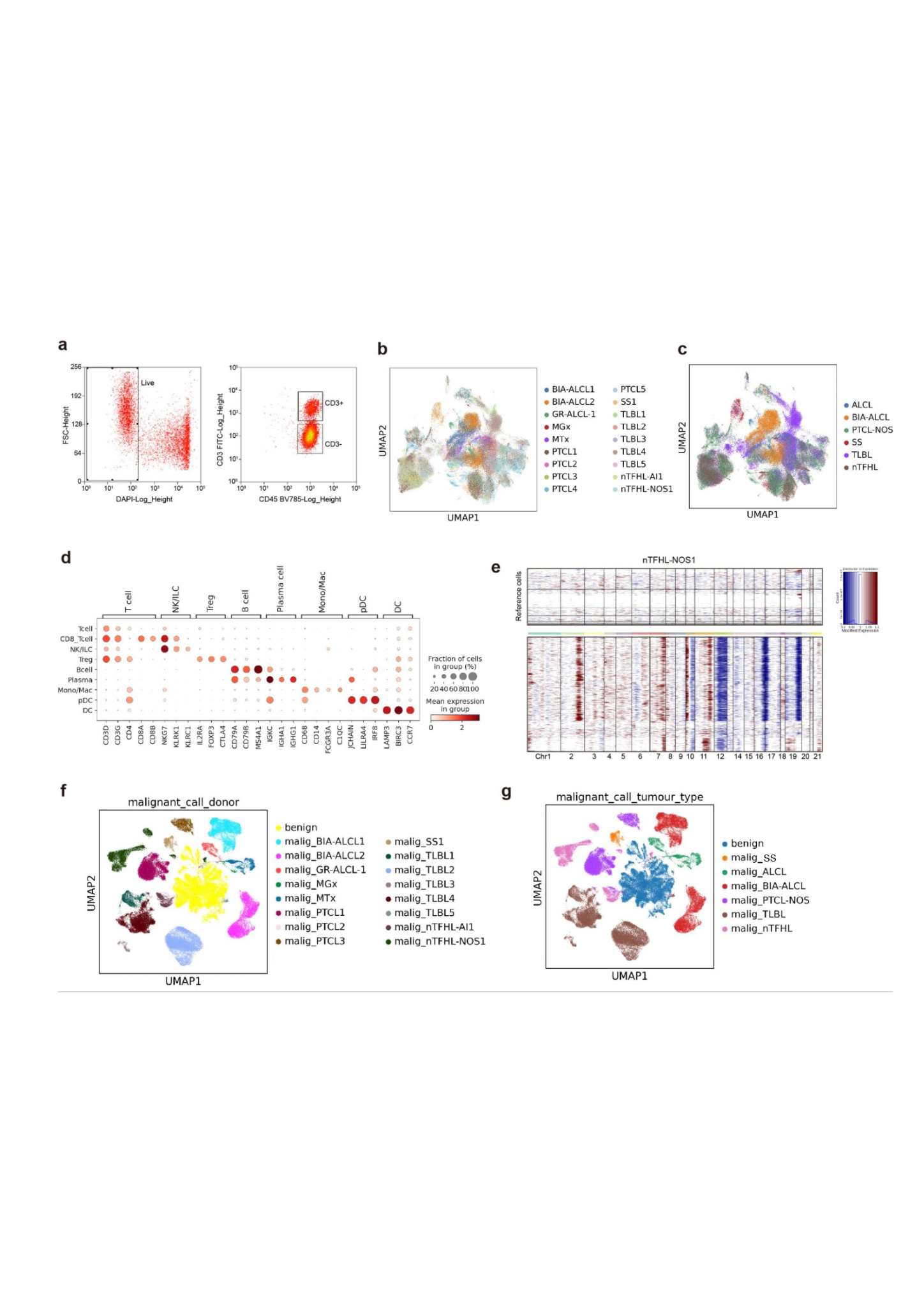

**Supplementary Figure 1. Overview of the TCL dataset.** (A) Representative gating strategy for FACS sorting of live CD3+/- CD45+ cells in suspension. DAPI staining positively identifies cells with compromised membrane integrity, which were excluded from the analysis. (B) A UMAP of the generated TCL dataset coloured according to donors. (C) A UMAP of the generated TCL dataset coloured according to disease type. (D) Dot plot showing the expression of marker genes for each cell type. (E) Heat map showing CNVs inferred from scRNA-seq data in a representative sample (nTFHL-NOS1). The top panel represents reference normal cells and the bottom heat map represents malignant T cells. (F) A UMAP of malignant and benign T cells in TCL coloured according to the donor origin. No malignant cells were identified in PTCL4 and PTCL5 cases with this analysis. (G) A UMAP of malignant and benign T cells in TCL coloured according to disease classification.

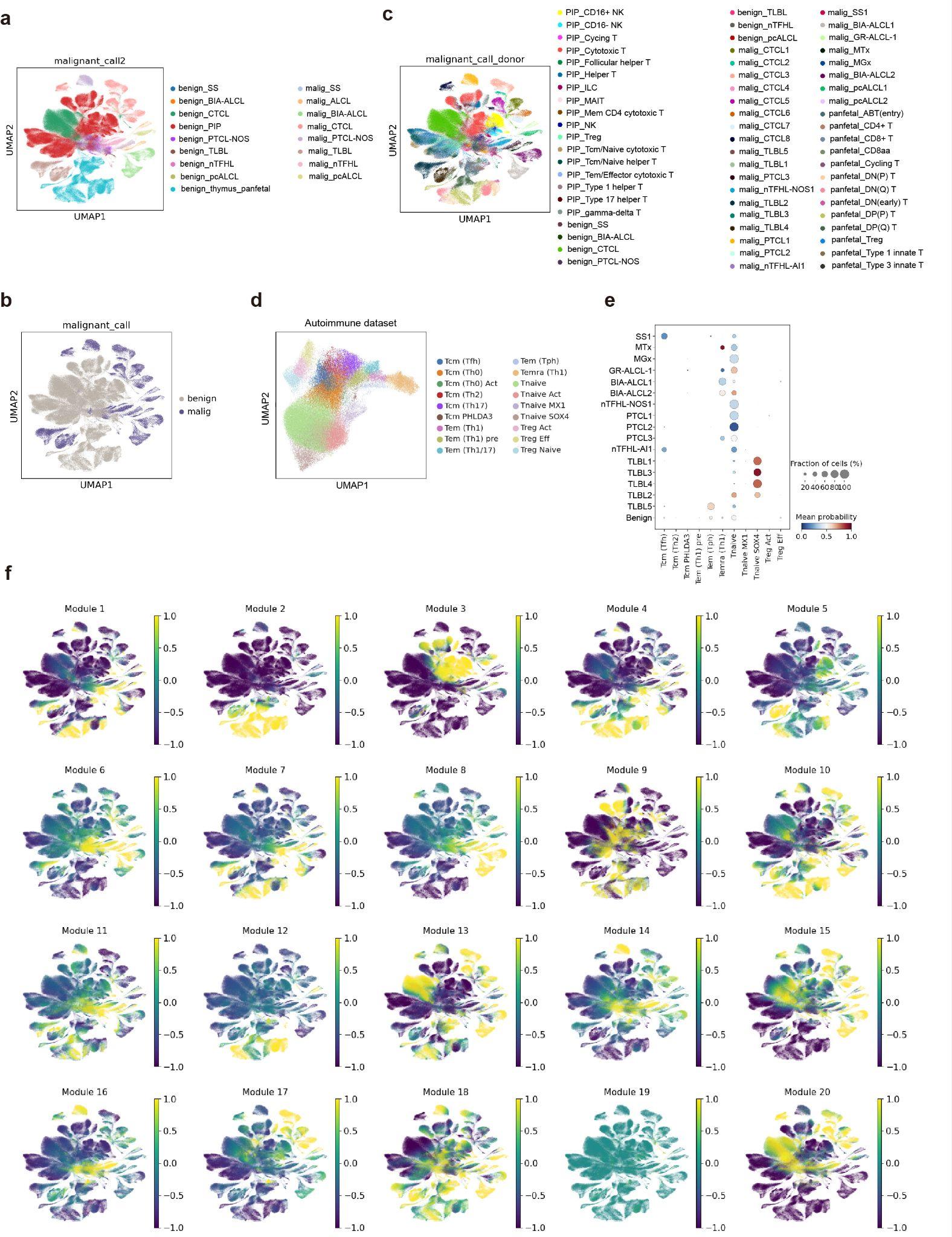

**Supplementary Figure 2. Integration of the current dataset with pan-body adult T cell, fetal T cell, CTCL, and pcALCL datasets.** (A) A UMAP of the integrated dataset coloured by sample type. (B) A UMAP showing benign and malignant T cells in the integrated dataset. (C) A UMAP of the integrated dataset coloured by cell type. (D) A UMAP projection of an autoimmunity T cell dataset. (E) CellTypist prediction of the cell of origin for each of the patient samples analysed, grouped according to histological entity and according to the autoimmune T cell atlas. (F) UMAPs showing expression of the 20 gene modules identified by Hotspot analysis.

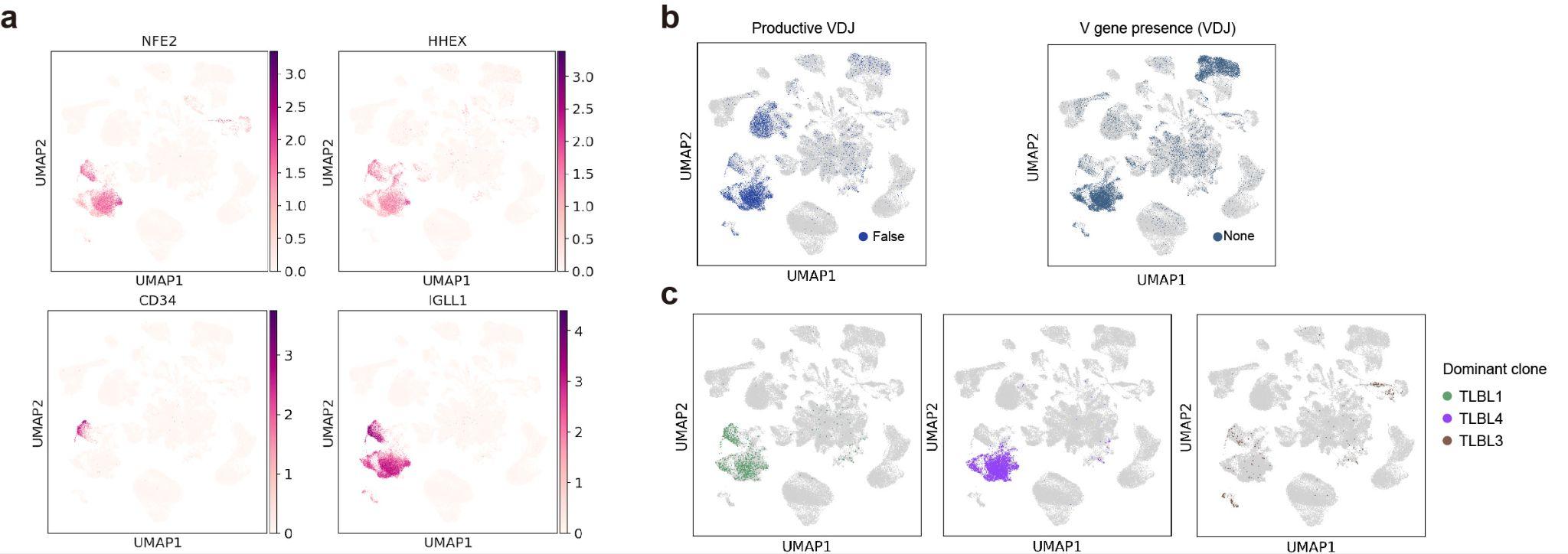

**Supplementary Figure 3. T cell populations in the dataset.** (A) Expression of immature T cell and haematopoietic stem/progenitor cell markers in TLBL. (B) UMAPs showing TCR contigs and V gene presence in TCL. (C) UMAPs showing the predominant TCR clonotype in each of the three TLBL cases analysed of a predicted double negative (DN) T cell cell of origin..

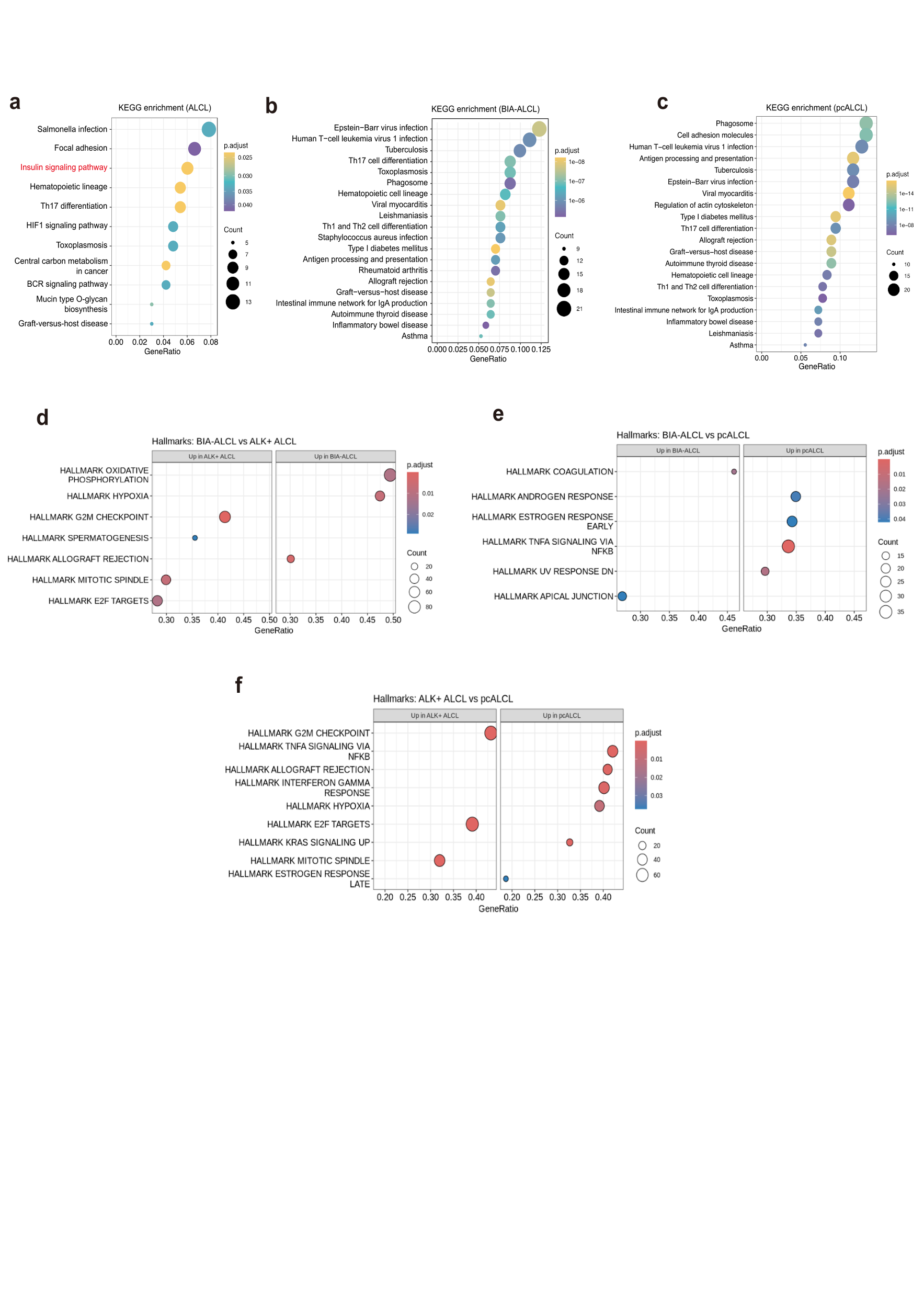

**Supplementary Fig 4. Pathway and Gene Set Enrichment Analysis of ALCL Subtypes.** KEGG pathway enrichment of upregulated genes in malignant T cells in ALK+ ALCL (A), BIA-ALCL (B) and pcALCL (C) compared to benign T cells. Gene set enrichment analysis (GSEA) was performed for comparisons of genetic programmes between BIA-ALCL and ALK+ ALCL (D), BIA-ALCL and pcALCL (E), and ALK+ ALCL and pcALCL (F). Samples were pseudo-bulked, and differential gene expression analysis was performed for each comparison of the indicated disease classifications. Genes were ranked by log fold change, and GSEA was conducted using the “Hallmark” gene sets from the Molecular Signatures Database (MSigDB). Dot plots display the top 10 enriched pathways by normalized enrichment score (NES), separated into up- and down-regulated pathways.

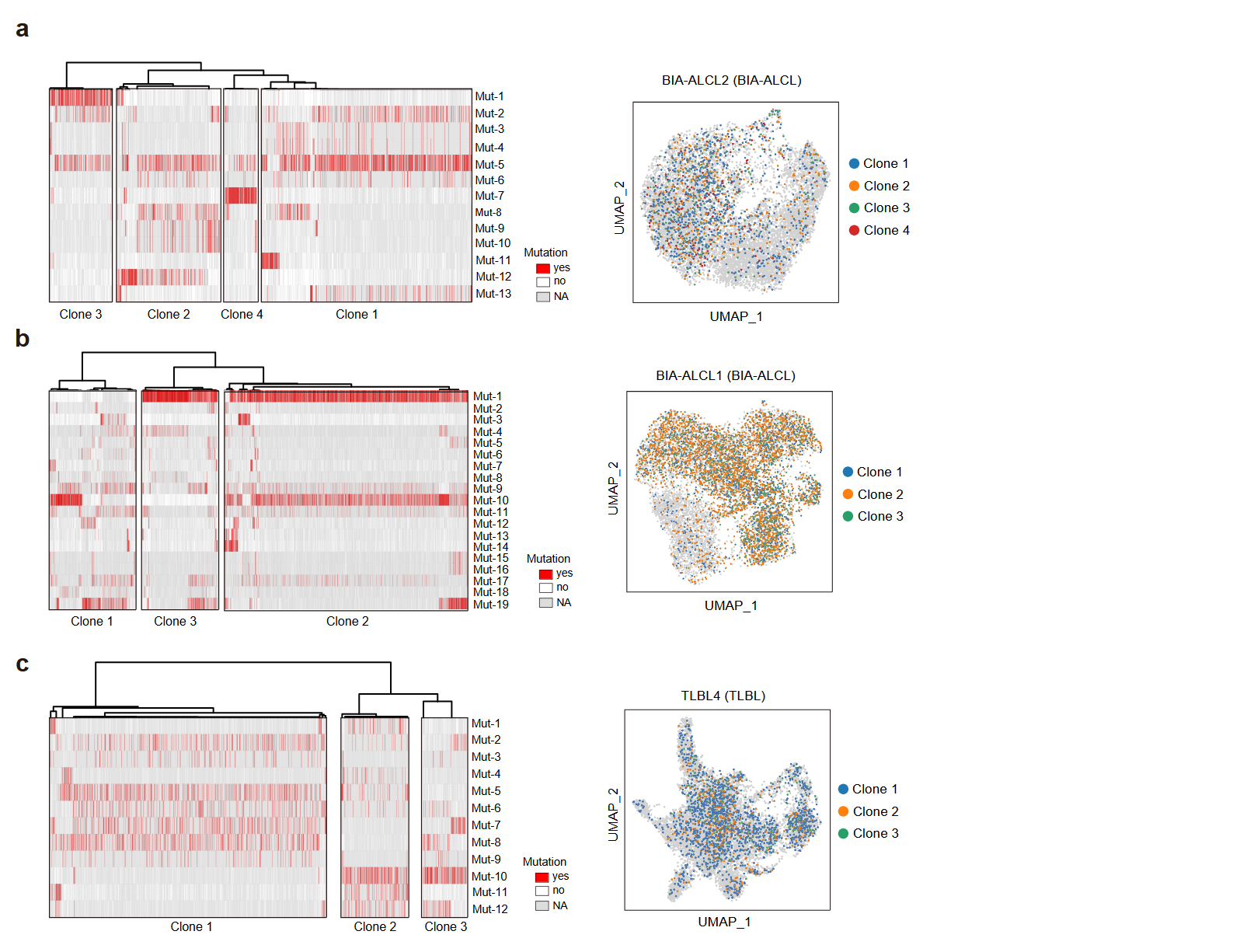

**Supplementary Figure 5. Intra-tumour genotypic and phenotypic heterogeneity.** (A-C) Left, heat maps of somatic mutations called from scRNA-seq data of representative tumours (BIA-ALCL2, BIA-ALCL1, and TLBL4). Right, UMAPs of malignant T cells from these three tumors coloured by genotypic clones. Grey represents unassigned clonal information.

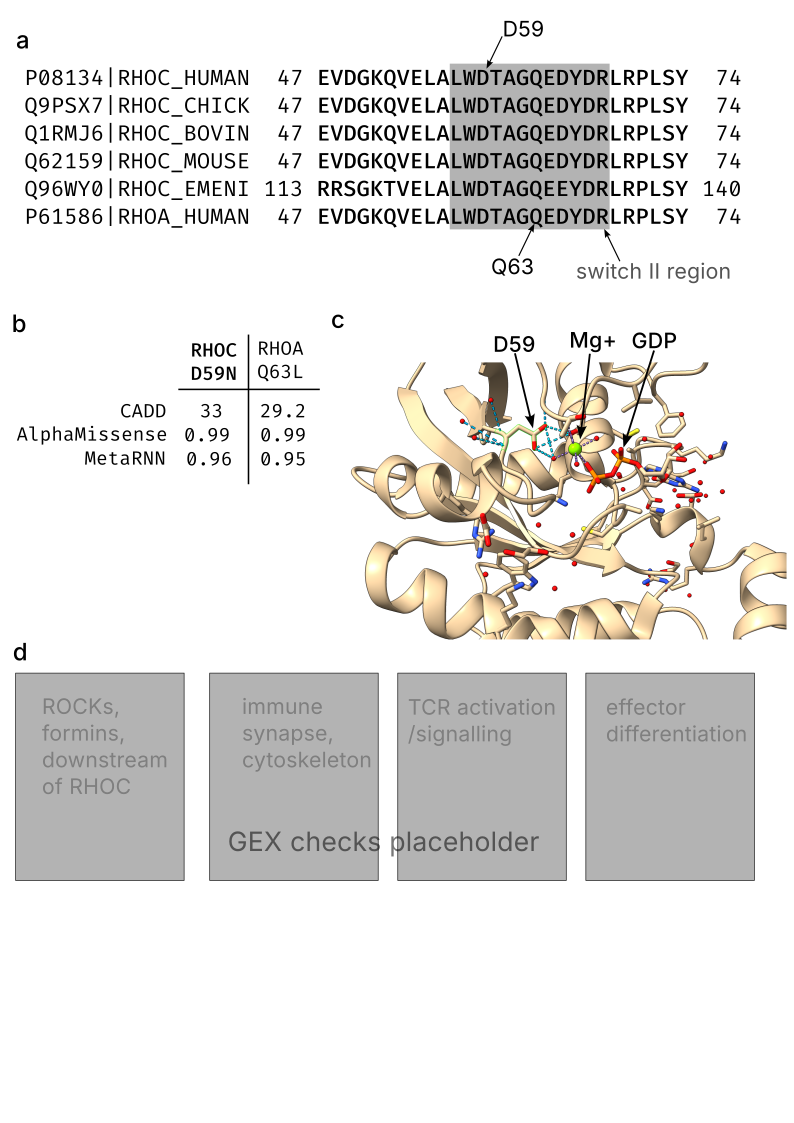
**Supplementary Figure 6. Predicted phenotypic impact of the RHOC mutation.** (A) Clustal Omega multiple species alignment (human, chicken, bovine, mouse, *Emericella nidulans* fungus) of RHOC and human RHOA demonstrating evolutionary conservation of sequence in the switch II region. The switch II region and positions of D59 and Q63 residues are highlighted. (B) Comparison of computational pathogenicity predictions for RHOC D59N and RHOA Q63L, which has been experimentally demonstrated to be activating^13^. (C) A ChimeraX render of the previously-published crystal structure of RHOC-GDP complex (2GCN) highlighting the D59 residue, Mg+ ion and GDP. Location of the mutated residue within the RHOC Mg+ binding pocket is shown.

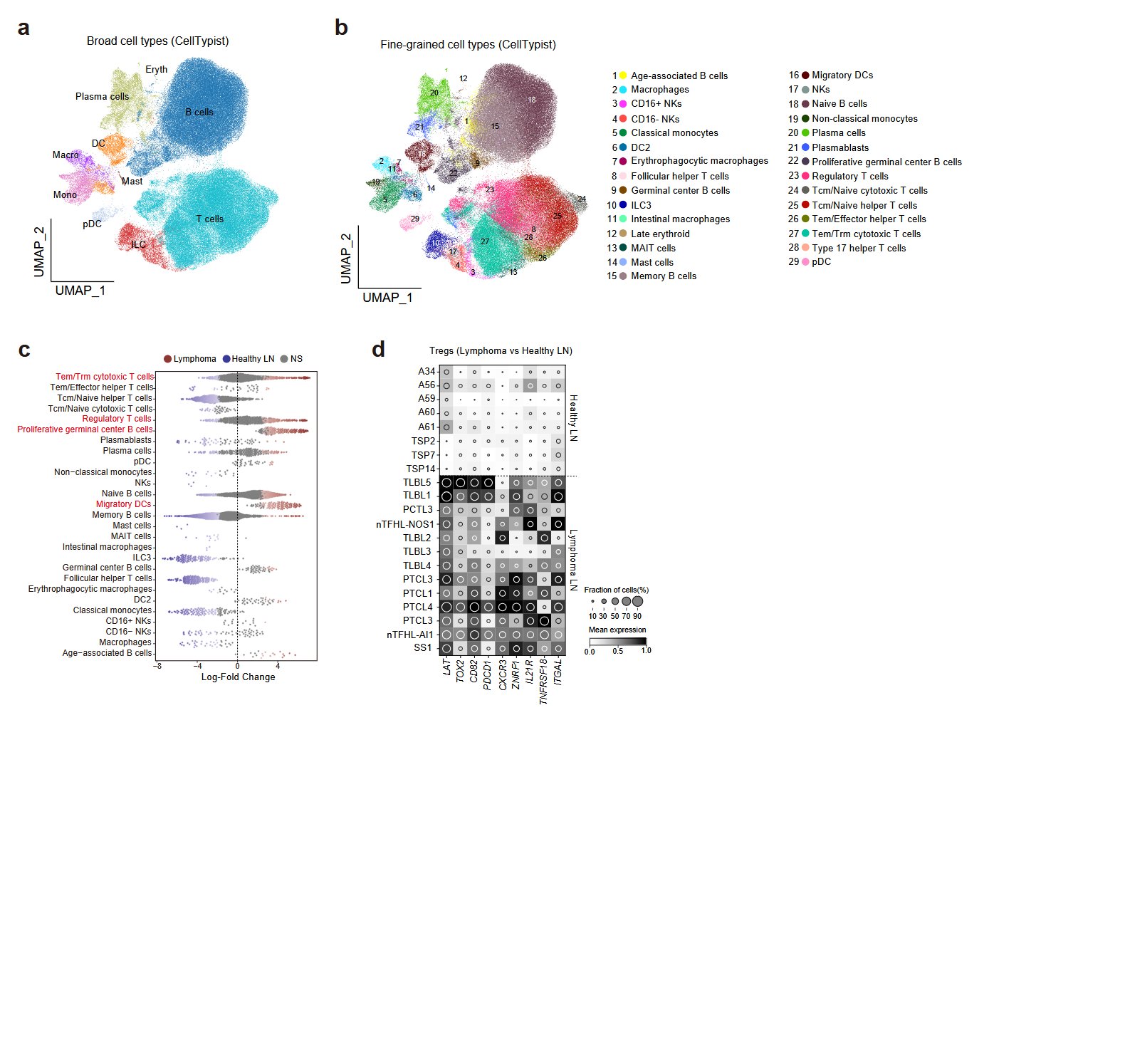

**Supplementary Figure 7. Comparison of TLBL, PTCL-NOS and nTFHL tumour lymph nodes with healthy lymph nodes from a pan-body atlas.** (A,B) Celltypist annotation of cell types (A) and subtypes (B). (C) Differential abundance of cell types. Red font indicates subtypes of interest. (D) Treg expression pattern across healthy and diseased lymph nodes. LN = lymph node; NS = not significant.

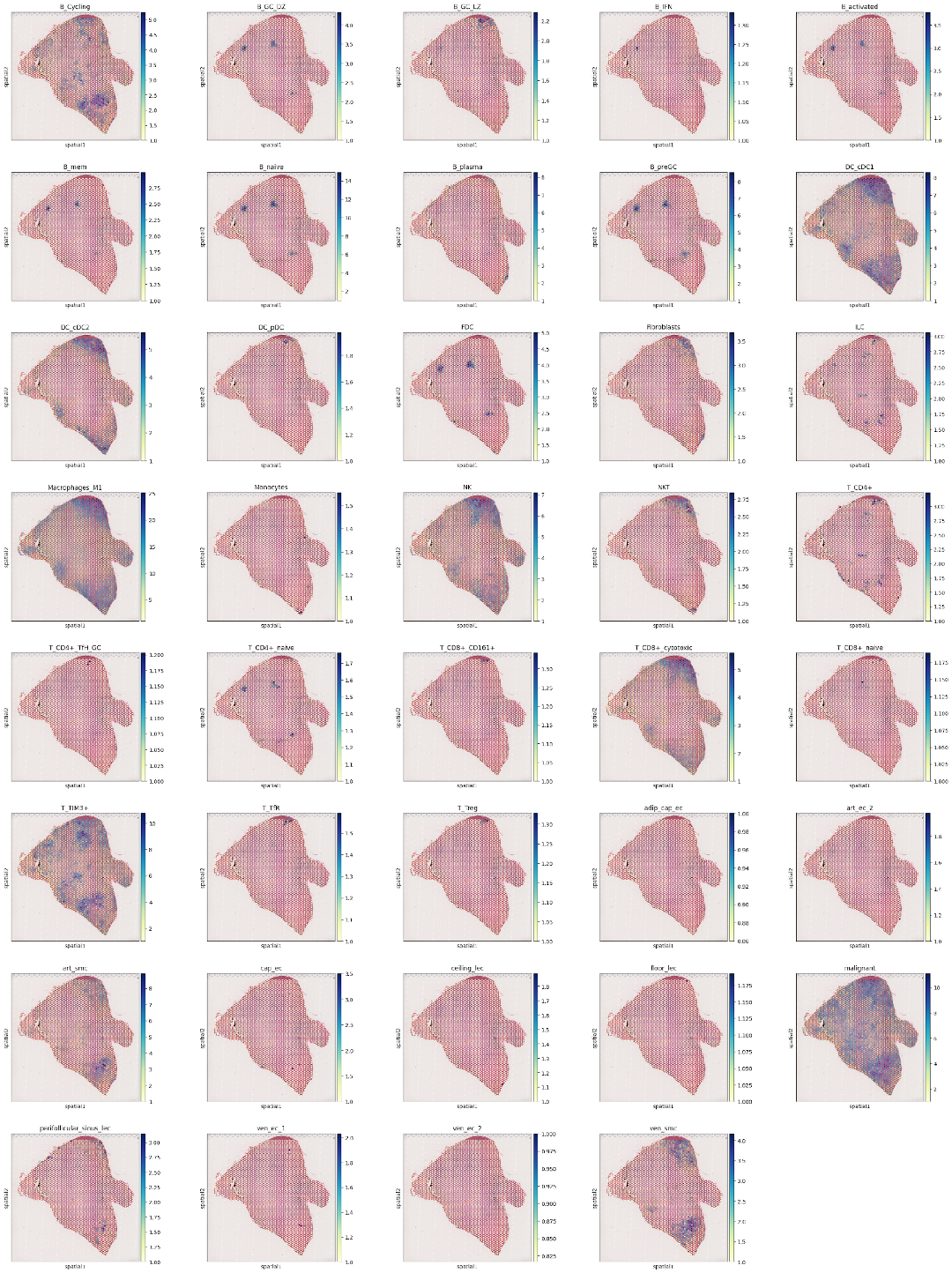

**Supplementary Figure 8. Cell2location predictions for all cell types in ALK+ ALCL Case 1.** Each subplot shows the predicted abundance of each cell type predicted by cell2location, using the 5% quantile of the posterior distribution to represent a confident lower boundary for cell abundance.

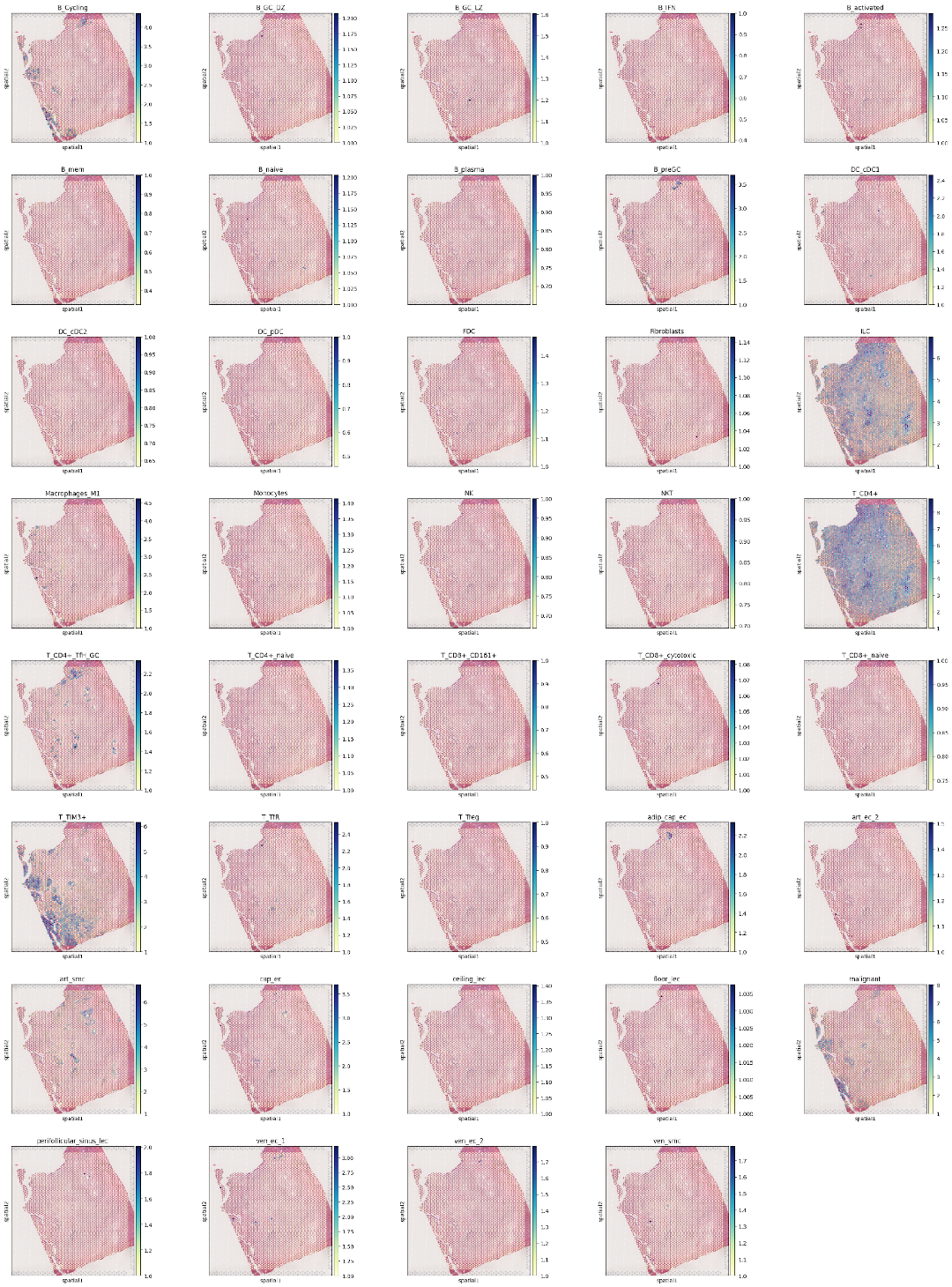

**Supplementary Figure 9. Cell2location predictions for all cell types in ALK+ ALCL Case 2.** Each subplot shows the predicted abundance of each cell type predicted by cell2location, using the 5% quantile of the posterior distribution to represent a confident lower boundary for cell abundance.

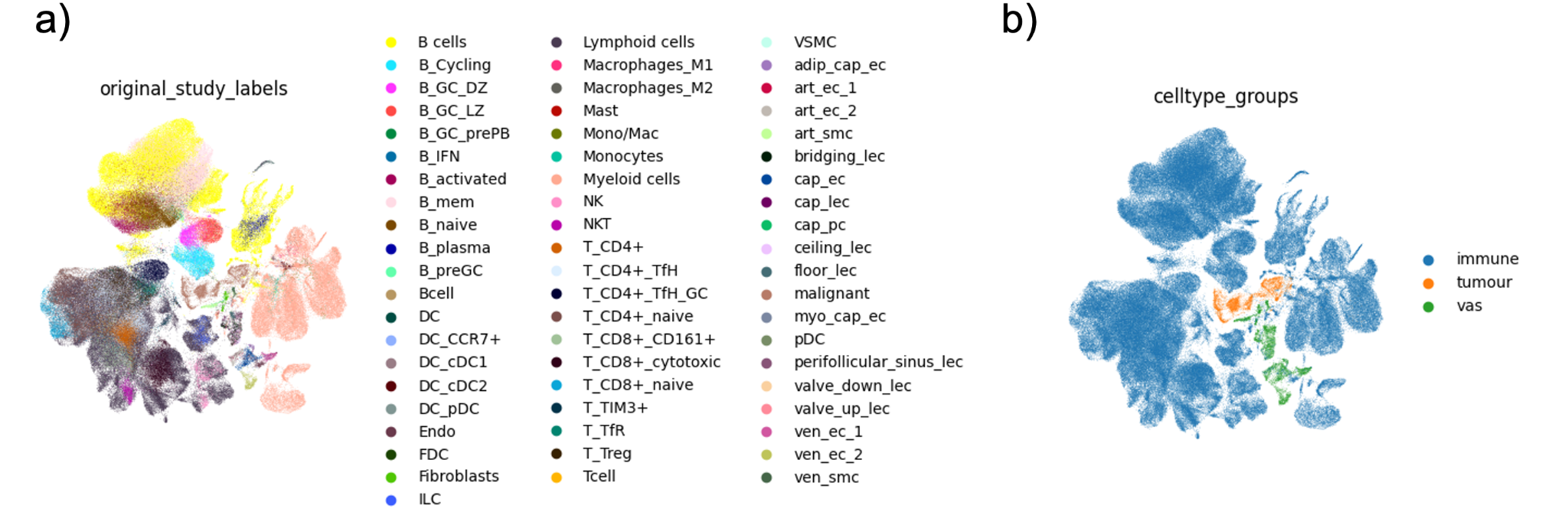

**Supplementary Figure 10. Integrated scRNAseq reference dataset for cell2location analysis.** (A) A UMAP representation showing the integrated scRNAseq datasets colored by original cell type labels. (B) A UMAP representation showing the integrated scRNAseq datasets colored by cell groups.

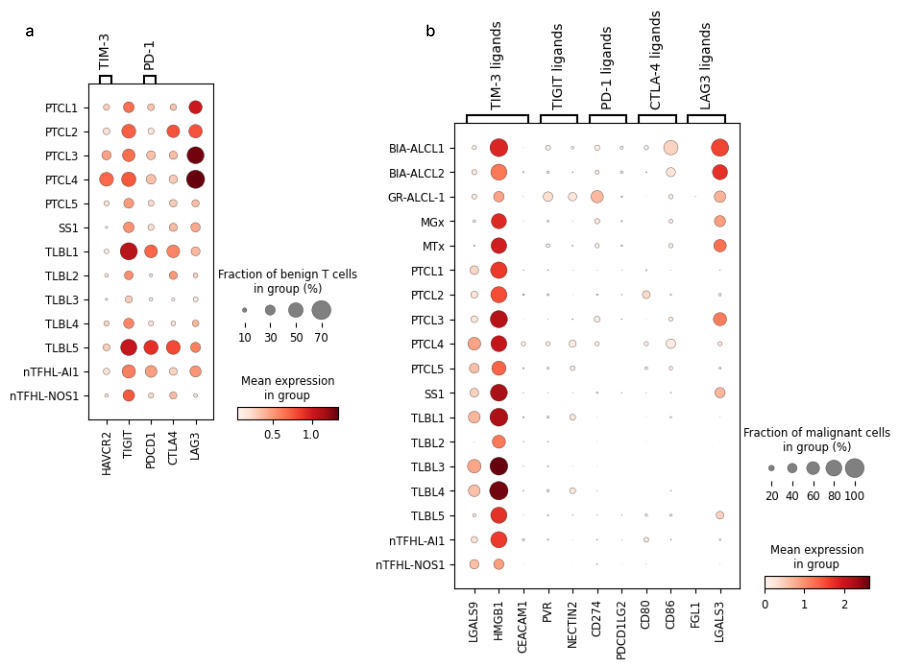
**Supplementary Figure 11. Expression of key immune checkpoint receptors and ligands in TCL.** (A) Dotplot of gene expression for immune checkpoint receptors in benign T cells from the cases listed, cases with < 100 benign T cells were excluded from this analysis. (B) Dotplot of gene expression for corresponding immune checkpoint receptor ligands in malignant cells of the indicated disease entities.

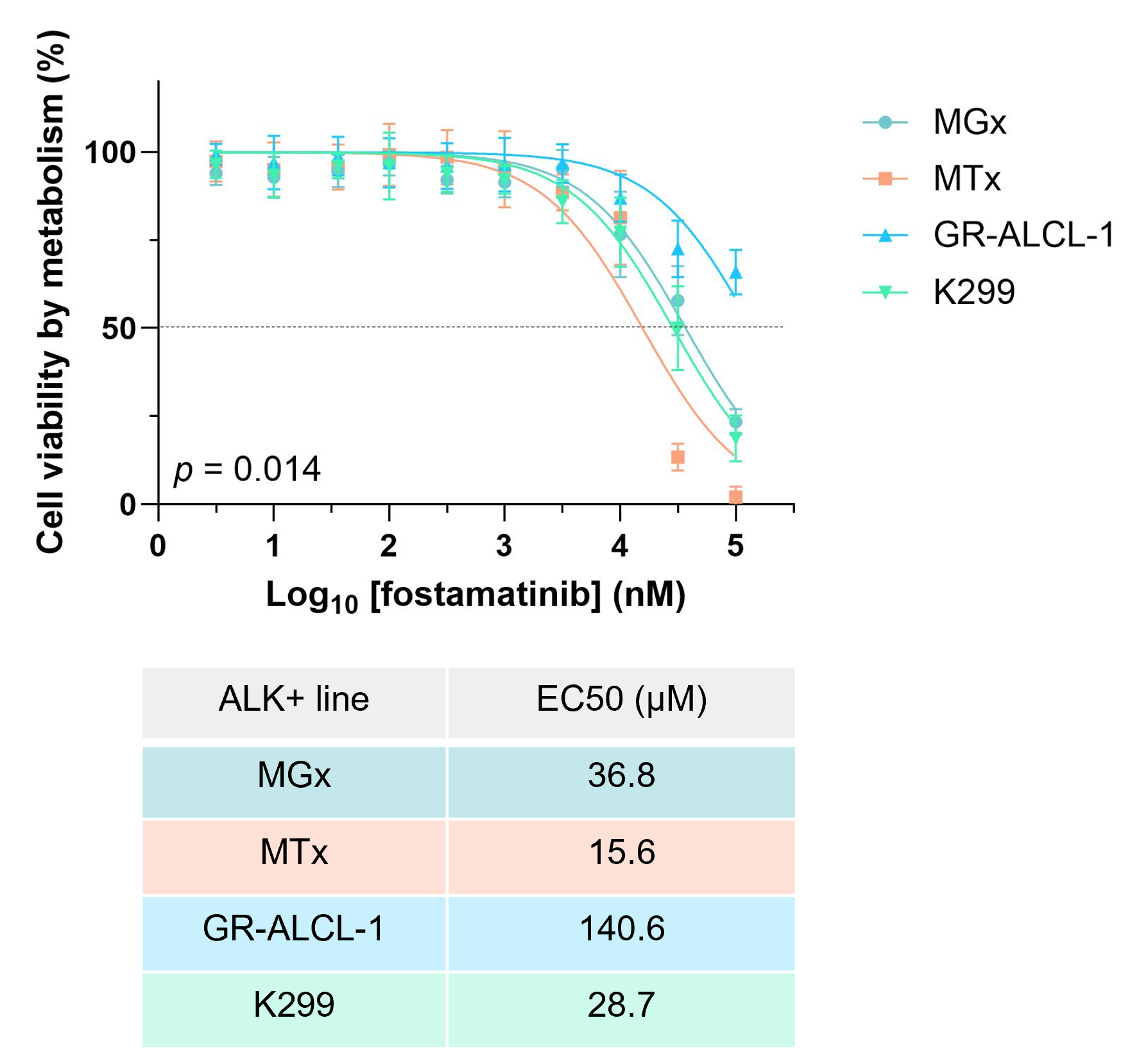

**Supplementary Figure 12. Validation of fostamatinib disodium as a potential efficacious therapy for the treatment of ALCL.** Ex vivo ALK+ ALCL PDX (MGx, MTx and GR-ALCL-1) and an established cell line (K299) cultured over 48 hours with the indicated doses of fostamatanib. EC50s were calculated using log(inhibitor) vs. normalized response. Data represent the means and standard deviations of biological triplicates, significance was calculated using ANOVA.
